# Defining the function of OmpA in the Rcs stress response

**DOI:** 10.1101/2020.07.15.203869

**Authors:** Juliette Létoquart, Kilian Dekoninck, Cédric Laguri, Pascal Demange, Robin Bevernaegie, Jean-Pierre Simorre, Olivia Dehu, Bogdan I. Iorga, Benjamin Elias, Seung-Hyun Cho, Jean-François Collet

## Abstract

OmpA, a protein commonly found in the outer membrane of Gram-negative bacteria, has served as a paradigm for the study of β-barrel proteins for several decades. In *Escherichia coli*, OmpA was previously reported to form complexes with RcsF, a surface-exposed lipoprotein that triggers the Rcs stress response when damage occurs in the outer membrane and the peptidoglycan. How OmpA interacts with RcsF and whether this interaction allows RcsF to reach the surface has remained unclear. Here, we integrated *in vivo* and *in vitro* approaches to establish that RcsF interacts with the C-terminal, periplasmic domain of OmpA, not with the N-terminal β-barrel, thus implying that RcsF does not reach the bacterial surface *via* OmpA. Our results reveal a novel function for OmpA in the cell envelope: OmpA competes with the inner membrane protein IgaA, the downstream Rcs component, for RcsF binding across the periplasm, thereby regulating the Rcs response.

## Introduction

The cell envelope is the morphological hallmark of Gram-negative bacteria. It consists of an inner membrane (IM) surrounding the cytoplasm as well as an outer membrane (OM), an asymmetric bilayer with phospholipids in the inner leaflet and lipopolysaccharides in the outer leaflet (Silhavy, Kahne, and Walker 2010). The two membranes are separated by the periplasm, a compartment in which lies a thin layer of peptidoglycan. The cell envelope is essential for viability: the OM serves as a permeability barrier against toxic compounds present in the environment while the peptidoglycan provides shape and osmotic protection to cells (Okuda et al. 2016, Typas et al. 2012, Egan, Errington, and Vollmer 2020).

Given the functional and structural importance of the envelope, bacteria need to respond to breaches in envelope integrity in a fast and adequate manner. Bacteria have therefore evolved sophisticated signaling systems that monitor envelope integrity and respond to perturbations (Ruiz and Silhavy 2005, MacRitchie et al. 2008, Delhaye, Collet, and Laloux 2019). In *Escherichia coli* and the Enterobacteriaceae, the Rcs system detects damage to the OM and the peptidoglycan (Wall, Majdalani, and Gottesman 2018, Laubacher and Ades 2008, Farris et al. 2010). In response, Rcs modulates the expression of dozens of genes, including those involved in the biosynthesis of colanic acid, an exopolysaccharide that accumulates on the cell surface to form a protective capsule (Wall, Majdalani, and Gottesman 2018, Laloux and Collet 2017).

Rcs signal transduction involves a multi-step phosphorelay (Wall, Majdalani, and Gottesman 2018). Under stress, the IM histidine kinase RcsC autophosphorylates, transfers the phosphoryl group to the IM protein RcsD and finally to the cytoplasmic response regulator RcsB. Remarkably, Rcs activity is modulated by two proteins that are not part of the phosphorylation cascade: RcsF and IgaA. RcsF is an OM lipoprotein that senses most Rcs-inducing cues, while IgaA is an essential IM protein that down-regulates Rcs (Takeda et al. 2001, Dominguez-Bernal et al. 2004) by interacting with RcsD (Wall, Majdalani, and Gottesman 2020). When perturbations occur in the peptidoglycan or in the OM, RcsF, while remaining anchored in the OM, reaches across the periplasm to interact with IgaA, leading this protein to alleviate its inhibition of the phosphorelay, turning on Rcs (Cho et al. 2014, Hussein et al. 2018). In the absence of stress, RcsF is occluded from IgaA by interacting with OM proteins. A complex between RcsF and BamA, the core component of the β-<u>b</u>arrel <u>a</u>ssembly <u>m</u>achinery (BAM), was identified (Cho et al. 2014, Konovalova et al. 2014) and its structure solved (Rodriguez-Alonso et al. 2020). This complex forms as an intermediate (Cho et al. 2014, Konovalova et al. 2014): delivery of unfolded OM β-barrels (OMPs) to BAM triggers the release of RcsF from BamA and its transfer to OMP partners (Rodriguez-Alonso et al. 2020). Three abundant OMPs (OmpA, OmpC and OmpF) have been identified as RcsF partners (Cho et al. 2014, Konovalova et al. 2014). Under stress conditions, newly synthesized RcsF molecules fail to interact with BamA (Cho et al. 2014): they remain in the periplasm, free to bind IgaA, triggering Rcs.

Crucially, whereas the general view is that OM lipoproteins are oriented toward the periplasm, previous work concluded that at least a portion of RcsF, a protein which is composed of an N-terminal disordered linker and a C-terminal globular domain required for signaling (Leverrier et al. 2011, Rogov et al. 2011), is exposed on the cell surface; OmpA, OmpC, and OmpF, but not BamA (Cho et al. 2014), were identified as potential vehicles for RcsF surface exposure (Cho et al. 2014, Konovalova et al. 2014, Konovalova, Mitchell, and Silhavy 2016). In a topological model of RcsF surface exposure (Konovalova et al. 2014), the lipid moiety of RcsF is anchored in the outer leaflet of the OM and the N-terminal disordered linker is exposed on the cell surface before being threaded through the lumen of the OMP partners. However, definitive evidence for this model is still lacking.

In addition, because OmpC, OmpF, and OmpA belong to two distinct structural groups, it is unclear whether RcsF interacts with its three OMP partners in a similar way. Indeed, OmpC and OmpF form 16-stranded β-barrels that associate into trimers in the OM. Because they form large β-barrels, they display a central pore (Basle et al. 2006, Yamashita et al. 2008, Radhakrishnan, Pritchard, and Viollier 2010, Housden et al. 2013) that is large enough to accommodate a disordered peptide such as the RcsF linker. The situation is less clear for OmpA: although this protein, the most abundant OMP in *E. coli*, has been studied for more than four decades, how it folds remains controversial. While some studies indicate that OmpA can also fold into a 16-stranded β-barrel with a large central pore (Singh et al. 2003, Stathopoulos 1996), the predominant view is that OmpA adopts a two-domain structure with an N-terminal 8-stranded β-barrel inserted in the OM (Pautsch and Schulz 1998) and a C-terminal, globular domain in the periplasm (De Mot and Vanderleyden 1994, Park et al. 2012). In this conformation, the β-barrel of OmpA is too small to accommodate a polypeptide. Thus, despite the tremendous work that has been done on OmpA, we do not know whether this protein adopts the large β-barrel structure when in complex with RcsF, or whether it folds into the predominant two-domain conformation. If the latter, where does RcsF bind OmpA?

To resolve these outstanding structural and mechanistic questions, here we dissected the OmpA-RcsF complex. By combining *in vivo* site-specific photo-crosslinking, targeted proteolysis, and nuclear magnetic resonance (NMR) titration, we established that OmpA adopts its two-domain structure when in complex with RcsF and that it is the C-terminal, periplasmic domain—not the β-barrel—that interacts with the lipoprotein. In addition, we identified residues in RcsF and in OmpA that are involved in the interaction, thus providing information about the binding interface. Taken together, our results indicate that the topology of OmpA-RcsF is different from that of OmpC/F-RcsF; they also imply that RcsF does not use OmpA to reach the cell surface. This has important implications for how RcsF senses OM stress: if the linker of RcsF is not on the surface in the OmpA-RcsF complex, then OmpA-RcsF cannot serve to monitor the state of the lipopolysaccharide leaflet via direct interactions with lipopolysaccharide molecules, as previously proposed (Konovalova, Mitchell, and Silhavy 2016). Finally, we determined the equilibrium dissociation constants of both the C-terminal domain of OmpA and the periplasmic domain of IgaA for RcsF and provide evidence that OmpA and IgaA compete for RcsF binding across the periplasm. Our results support a model in which OmpA serves as a buffer for RcsF, titrating it from IgaA, thereby fine-tuning Rcs activity.

## Results

### RcsF interacts with the C-terminal region of OmpA *in vivo*

The stress sensor lipoprotein RcsF was previously shown to be surface-exposed (Cho et al. 2014, Konovalova et al. 2014), and OmpA was described as a possible vehicle for its surface exposure (Cho et al. 2014, Konovalova et al. 2014). However, how OmpA folds when in complex with RcsF and whether this interaction allows RcsF to become surface-exposed remain to be determined.

To close this gap, we characterized the OmpA-RcsF interaction. In a previous study, we identified six RcsF residues (in the N-terminal disordered linker and at the tip of the signaling domain) as being part of the interaction interface between RcsF and OmpA (Cho et al. 2014). These residues were identified using a site-specific photo-crosslinking strategy in which a photoreactive, crosslinkable amino acid is inserted at specific positions in the protein of interest, with the help of an exogenous orthogonal tRNA/aminoacyl-tRNA synthetase pair (Chin et al. 2002). We first sought to confirm and extend these results to more clearly define the binding interface in RcsF. Instead of using the hydrophobic crosslinker *p*-benzoyl-L-phenylalanine like before (Cho et al. 2014), we used *N*^6^-((3-(3-methyl-3*H*-diazirin-3-yl)propyl)carbamoyl)-*L*-lysine (DiZPK), a lysine analog with substantially higher photo-crosslinking efficiency than *p*-benzoyl-L-phenylalanine (Zhang et al. 2011). We selected 11 positions distributed along the RcsF sequence, including four (R21, Q28, Q33, R45) in the disordered linker and seven on the surface of the signaling domain (N54, Q79, R89, K98, E110, P116, Q121) (Figure 1). After UV illumination, a ~55 kDa band, the size of the OmpA (40 kDa)-RcsF (14 kDa) complex, formed and was detected with an anti-RcsF antibody (Figure 1) for the following variants (in decreasing intensity): RcsF_R89X_, RcsF_P116X_, RcsF_Q79X_, RcsF_R45X_, RcsF_K98X_, RcsF_Q121X_, RcsF_N54X_, and RcsF_E110X_. The identity of the ~55 kDa band as OmpA-RcsF was further verified by showing that it did not form in Δ*ompA* cells (Figure 1 – figure supplements 1 and 2). Thus, as in our previous study (Cho et al. 2014), complex formation was observed when the photoactivatable amino acid was inserted in the linker (R45) or at the tip of the signaling domain (Q79 and P116). The complex also formed when DiZPK was incorporated in other regions of the signaling domain, such as a-helix 1 (N54), a-helix 2 (R89 and K98), β-strand 2 (E110), and β-strand 3 (Q121) (Figure 1). Taken together, these observations substantially enlarge the region of RcsF known to interact with OmpA. Of note, we observed that the RcsF_R45X_, RcsF_R89X_ and RcsF_K98X_ variants formed a UV-dependent band migrating slightly higher than the OmpA-RcsF complex (Figure 1). Focusing on RcsFR89X, we identified this band as a complex between RcsF and OmpC/F because it did not form when *ompR*, a transcription factor required for the production of OmpC and OmpF (Hall and Silhavy 1979, Chubiz and Rao 2011), was deleted (Figure 1 – figure supplement 2).

**Figure 1.**
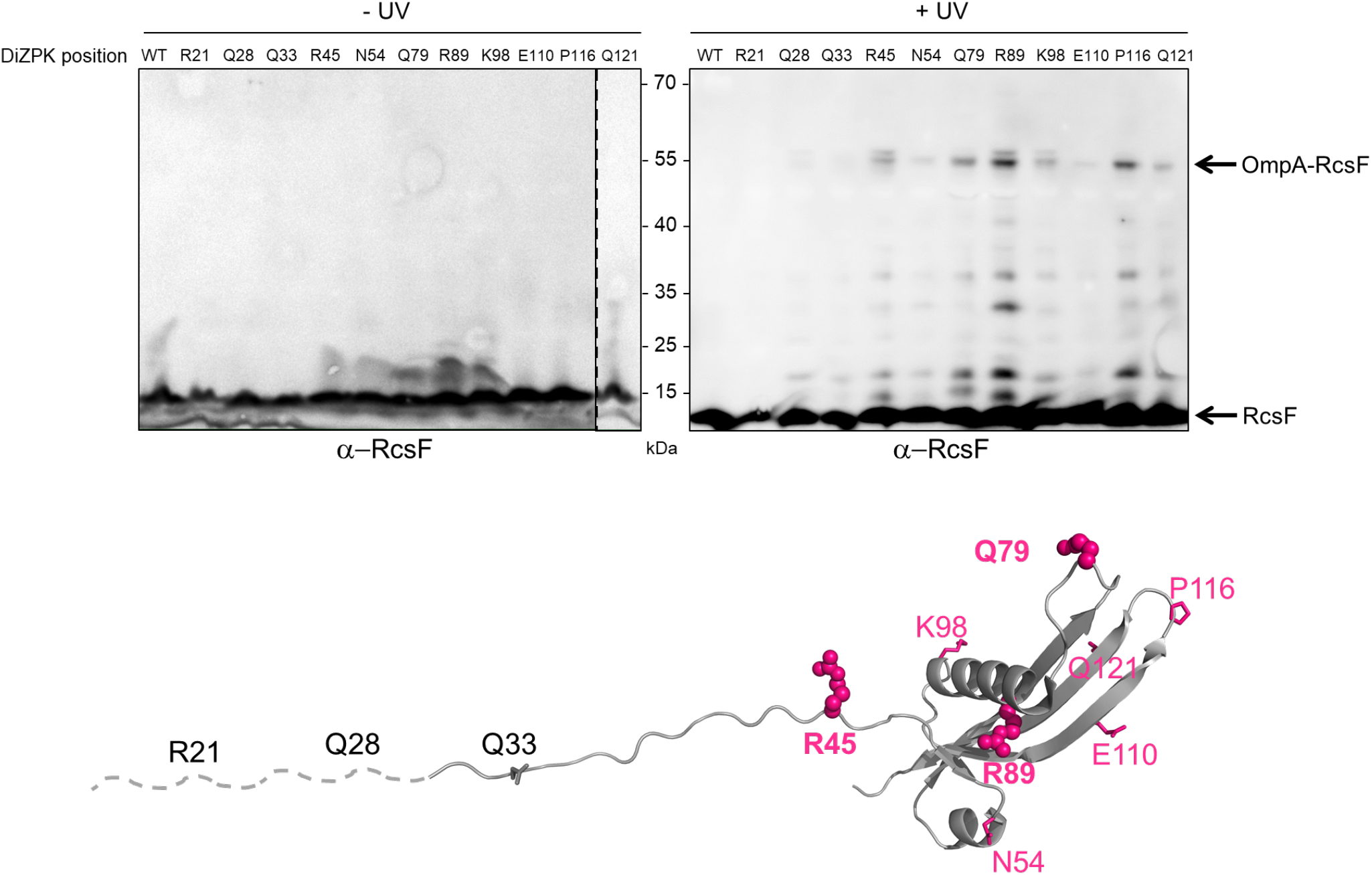
Defining the binding interface of RcsF on OmpA using *in vivo* site-specific photo-crosslinking. Upper panel: *ΔrcsF* cells expressing wild-type (WT) RcsF or DiZPK-containing RcsF variants (from pSC253) were irradiated with UV light (+) or not (-), and protein samples were immunoblotted with an anti-RcsF antibody. A 55 kDa band, corresponding to the size of the OmpA-RcsF complex, was observed for eight of the mutants (R45, N54, Q79, R89, K98, E110, P116, Q121). Lower panel: residues of RcsF were replaced by DiZPK to map the zone of interaction with OmpA. In this cartoon of the NMR structure of RcsF (PDB: 2L8Y), the truncated N-terminal portion of the protein is shown as a dashed line and the residues that were found to interact with OmpA appear in magenta. The side chains of the residues that were selected for further experiments are shown as spheres, and other side chains are represented as sticks.

We next sought to identify where RcsF binds OmpA. The general view is that OmpA consists of two domains, an 8-stranded β-barrel anchoring the protein in the OM and a soluble C-terminal domain located in the periplasm, where it binds the peptidoglycan (De Mot and Vanderleyden 1994, Park et al. 2012) (Figure 2A). However, an alternative conformation has been proposed in which OmpA folds into a large, 16-stranded β-barrel (Singh et al. 2003, Stathopoulos 1996) (Figure 2A). To gain insight into where RcsF binds OmpA and to characterize OmpA’s conformation when in complex with RcsF, we first engineered an OmpA variant with a thrombin cleavage site inserted after residue V189 (OmpA_TH_189_), in the middle of the OmpA sequence (Figure 2B). Taking the two-domain structure as a reference, the cleavage site was inserted between the N- and C-terminal domains. We then selected three DiZPK-containing RcsF mutants (RcsFR45X, RcsFQ79X, RcsFR89X) that formed a covalent complex with OmpA at high levels when exposed to UV light (Figure 1). These variants display DiZPK in three regions of RcsF: in RcsFR45X, DiZPK is present at the end of the disordered linker, while RcsF_Q79X_ displays DiZPK in the large loop at the tip of the signaling domain and RcsF_R89X_ displays it on the central a-helix 2 (Figure 1). These variants were expressed in *E. coli* cells also producing OmpA_TH_189_, and complex formation was induced with UV light (Figure 2C). For all three RcsF variants, digestion of the 55-kDa OmpA_TH_189_-RcsF complex with thrombin yielded a ~35 kDa band that was recognized by both anti-RcsF and anti-His antibodies (Figure 2C, D). Given the presence of a His-tag in the C-terminus of OmpA (Materials and Methods), we concluded that RcsF (14 kDa) interacts with the C-terminal region of OmpA (~16 kDa) *in vivo*.

**Figure 2.**
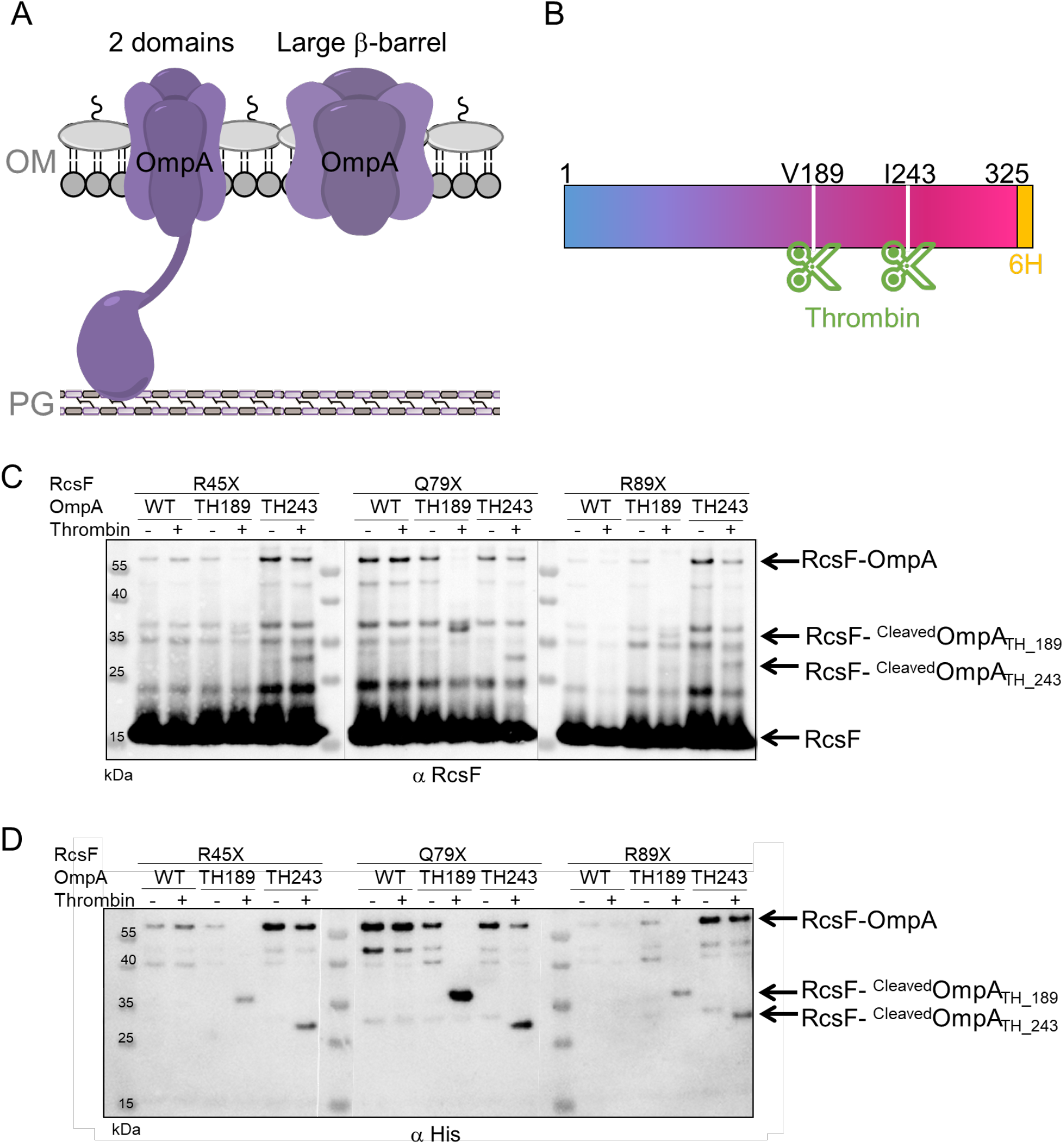
RcsF interacts with the C-terminal portion of OmpA *in vivo.* (A) Schematic of the two conformations of OmpA. Left: the predominant view is that OmpA adopts a two-domain structure, with an N-terminal β-barrel embedded in the OM and a C-terminal periplasmic domain binding the peptidoglycan. Right: an alternative conformation in which OmpA folds into a large β-barrel has also been proposed (Singh et al. 2003, Stathopoulos 1996). (B) Schematic of OmpA variants containing a thrombin-specific cleavage site (scissors at positions V189 and I243) and a 6x-histidine tag (orange) at the C-terminus. (C-D) *In vivo* site-specific photo-crosslinking of RcsF. *ΔrcsF* cells co-expressing one of the DiZPK-containing RcsF variant (R45X, Q79X, or R89X) together with OmpA (wild-type, or with a thrombin site inserted at V189 or I243) were UV-irradiated. The RcsF variants were expressed from pSC253; OmpA, OmpATH_189 and OmpATH_243 were expressed from the chromosome. After immunoprecipitation with anti-RcsF, protein samples were incubated (+) or not (-) with thrombin and immunoblotted with an anti-RcsF (C) or an anti-His-tag antibody (D). At least partial digestion of the ~55 kDa complex corresponding to OmpA-RcsF occurred with all three DiZPK-containing RcsF variants, yielding a band (RcsF-^Cleaved^OmpA_TH_189_ or RcsF-^cleaved^ OmpA_TH_243_) migrating at lower molecular weights that was detected by both antibodies.

To further delineate the region of OmpA that is important for complex formation, a second OmpA mutant with a thrombin cleavage site inserted after residue I243 (OmpA_TH_243_) was generated (Figure 2B). Like OmpA_TH_189_, OmpA_TH_243_ (~10 kDa) could be photo-crosslinked to the three DiZPK-containing RcsF variants described above (Figure 2C, D). In all three cases, thrombin digestion of the OmpA_TH_243_-RcsF complexes generated a smaller band (~30 kDa) that was recognized by anti-RcsF antibodies (Figure 2C, D). In contrast to OmpA_TH_189_-RcsF, which underwent complete digestion, OmpA_TH_243_-RcsF only underwent partial cleavage (Figure 2C, D), which probably reflected decreased accessibility of the cleavage site to the protease. This ~30 kDa band was also detected by anti-His antibodies (Figure 2D), indicating that the three tested residues of RcsF bind the region of OmpA between residue I243 and the C-terminus.

### The C-terminal region of OmpA is necessary and sufficient for binding RcsF

The results above indicated that RcsF interacts with the C-terminal region of OmpA, raising the question of whether the N-terminal region also participates in this interaction. To probe this potential interaction directly, we generated an OmpA variant lacking the C-terminal moiety (OmpA_1-170_) and tested whether it could be crosslinked to RcsF (Figure 3A). Chemical crosslinking was carried out with 3,3’-dithio-bis[sulfosuccinimidylpropionate] (DTSSP) (Cho et al. 2014). Whereas complexes formed in cells expressing wild-type OmpA, no complex was detected in cells producing OmpA_1-170_ (Figure 3A). In addition, expression of OmpA_1-170_ did not suppress the activation of the Rcs system (Figure 3A) that occurs in cells lacking OmpA and that results from the inability of RcsF to interact with this β-barrel (Cho et al. 2014).

**Figure 3.**
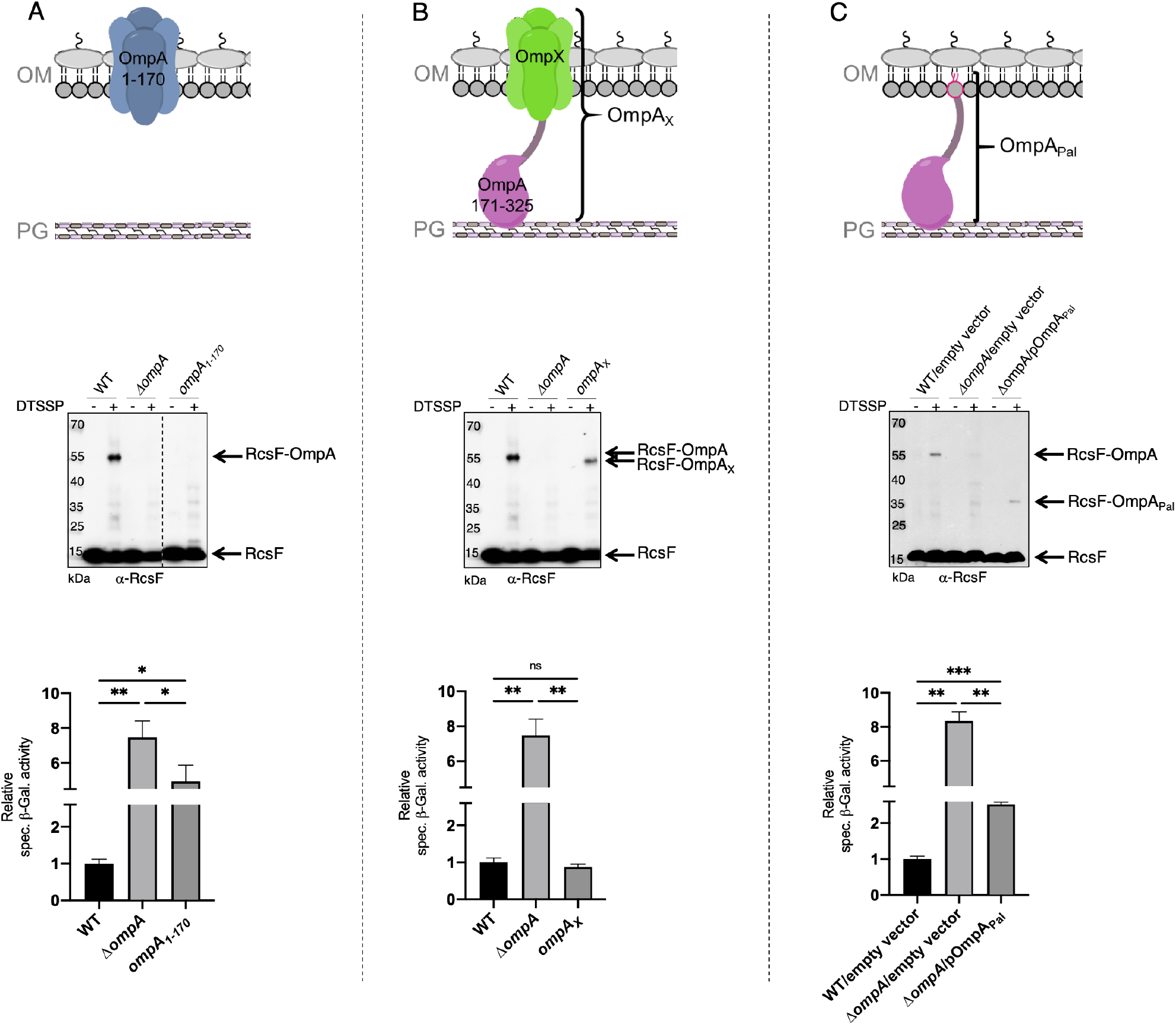
The periplasmic domain of OmpA is necessary and sufficient for the interaction with RcsF *in vivo.* (**A**) *Upper panel:* Schematic of the truncated OmpA variant corresponding to the 8-stranded β-barrel domain (OmpA_1-170_). *Middle panel: In vivo* chemical crosslinking of RcsF to OmpA and OmpA_1-170_. WT, *ΔompA* and *ΔompA::ompA_1-170_* DH300 cells were incubated with or without 3,3’-dithio-bis[sulfosuccinimidylpropionate] (DTSSP). Proteins were immunoblotted with anti-RcsF (same for the middle panel in **B** and **C**). The OmpA-RcsF complex was detected in WT cells but not in cells expressing OmpA_1-170_. *Lower panel:* β-galactosidase (β-gal) activity was measured using the transcriptional *rprA-lacZ* fusion on the chromosome (Majdalani, Hernandez, and Gottesman 2002) using the same DH300 strains as in the middle panel (same for the lower panel in **B** and **C**). Deleting *ompA* induces Rcs, expression of OmpA_1-170_ does not restore basal Rcs activity. (**B**) *Upper panel:* Schematic of the hybrid protein consisting of a fusion between the 8-stranded β-barrel OmpX and the periplasmic domain of OmpA (OmpA_171-325_) (OmpA_X_). *Middle panel: In vivo* chemical crosslinking of RcsF to OmpA and OmpA_X_. WT, *ΔompA* and *ΔompA::ompA_X_* cells were incubated with or without DTSSP. The RcsF-OmpA complex was detected in WT cells as well as in cells expressing OmpA_X_ (RcsF-OmpA_X_). *Lower panel:* Deleting *ompA* induces Rcs, expression of OmpA_X_ restores basal Rcs activity. (**C**) *Upper panel:* Schematic of the C-terminal domain of OmpA (OmpA_171-325_) fused to the signal sequence and lipobox of the OM lipoprotein Pal (OmpAPal). *Middle panel: In vivo* chemical crosslinking of RcsF to OmpA and OmpA_Pal_. WT and *ΔompA* harboring pDSW204, an empty vector, used as control, and *ΔompA* cells harboring pKiD22, expressing OmpA_Pal_ from an IPTG-inducible promoter, were incubated with or without DTSSP. The OmpA-RcsF complex was detected in WT cells as well as in cells expressing OmpAPal (OmpAPal-RcsF). *Lower panel:* Deleting *ompA* induces Rcs, expression of OmpA_Pal_ substantially decreases Rcs activity. Mean (n=3) and standard deviation (error bars) are shown. Differences were evaluated with Student’s *t* test (ns, not significant; *p<0.05; ***p<0.001).

To test whether the N-terminal domain of OmpA was indirectly required for the OmpA-RcsF complex to form, we generated a hybrid protein (OmpA_X_) in which the C-terminal region of OmpA (OmpA_171-325_), corresponding to the periplasmic domain in the two-domain structure, was fused to OmpX (Figure 3B). OmpX is a small 8-stranded β-barrel that constitutes a structural homolog of the N-terminal region of OmpA when it folds as a small β-barrel (the two β-barrels can be superimposed with a root mean square deviation (RMSD) of 2.49 Å; Figure 3 – figure supplement 1A). OmpX does not share sequence homology with the N-terminal region of OmpA (~25%) and can thus be considered as an OM anchor for the C-terminal region when fused to the latter as in OmpA_X_ (Figure 3B). We found that OmpA_X_ could be crosslinked to RcsF (Figure 3B) and that its expression fully suppressed Rcs activation (Figure 3B). Because we could not completely exclude the unlikely possibility that OmpAX could rearrange into a large β-barrel able to bind RcsF, we prepared an additional variant of OmpA (OmpAPal) in which the C-terminal domain (OmpA_171-325_) was fused to the signal sequence and lipobox (for lipid modification; (Szewczyk and Collet 2016)) of the OM lipoprotein Pal, thus converting the C-terminal domain of OmpA into a lipoprotein (Figure 3C). Remarkably, expression of OmpA_Pal_ led to the formation of OmpA_Pal_-RcsF and substantially decreased Rcs activity (Figure 3C). In these experiments, we confirmed that the expression levels of OmpA_1-170_, OmpA_X_ and OmpA_Pal_ were similar to those of the wild type (Figure 3 – figure supplement 2). Thus, the C-terminal region of OmpA is necessary and sufficient for the interaction with RcsF, and the N-terminal region is dispensable. Altogether, these results are consistent with the conclusion that OmpA adopts its two-domain structure—and not the large β-barrel conformer—when in complex with RcsF. We therefore conclude that RcsF interacts with the C-terminal, globular domain of OmpA and that this interaction takes place on the periplasmic side of the OM.

### RcsF interacts with the periplasmic domain of OmpA *in vitro*

We next sought to validate our *in vivo* observations by probing the formation of a complex between RcsF and the soluble, periplasmic domain of OmpA (OmpA_186-325_) *in vitro* using purified proteins. However, attempts to pull-down OmpA_186-325_ with a soluble, His-tagged version of RcsF failed (data not shown), suggesting that the interaction between these two proteins was weak. Because NMR is a highly effective tool to investigate weak protein-protein interactions (Vaynberg and Qin 2006), we employed NMR titration experiments of ^15^N-labeled OmpA_186-325_ by RcsF. In this approach, the ^15^N-^1^H 2D NMR spectra of OmpA were recorded upon addition of increasing amounts of RcsF (2 and 10 molar equivalents); when OmpA and RcsF interact, concentration-dependent perturbations in the NMR spectra appear. Remarkably, upon addition of RcsF, several OmpA residues showed chemical shift variations in ^15^N-^1^H 2D correlation experiments (Figure 4A, B). Most of the shifted residues (T240, G244, S245, D246, A247, G251, L252, K294 and A297; Figure 4B) were located near the tip of the periplasmic domain (Figure 4C, D), identifying this region, and in particular a flexible loop between β-strand 2 and a-helix 3, as part of the binding interface with RcsF. The importance of this loop for the OmpA-RcsF interaction was confirmed using site-specific photo-crosslinking: OmpA was strongly crosslinked to RcsF when DiZPK was introduced at residue D246, while weak complex formation was observed with OmpA_R242X_ and OmpA_Y248X_ (Figure 4E). It is remarkable that these results nicely fit with those obtained using OmpATH_243 (Figure 2B) that identified the same region of the C-terminal domain of OmpA as part of the zone of interaction with RcsF. Thus, taken together, our results allow us to conclude that RcsF interacts with the C-terminal domain of OmpA in its globular conformation, not only *in vivo* but also *in vitro*.

**Figure 4.**
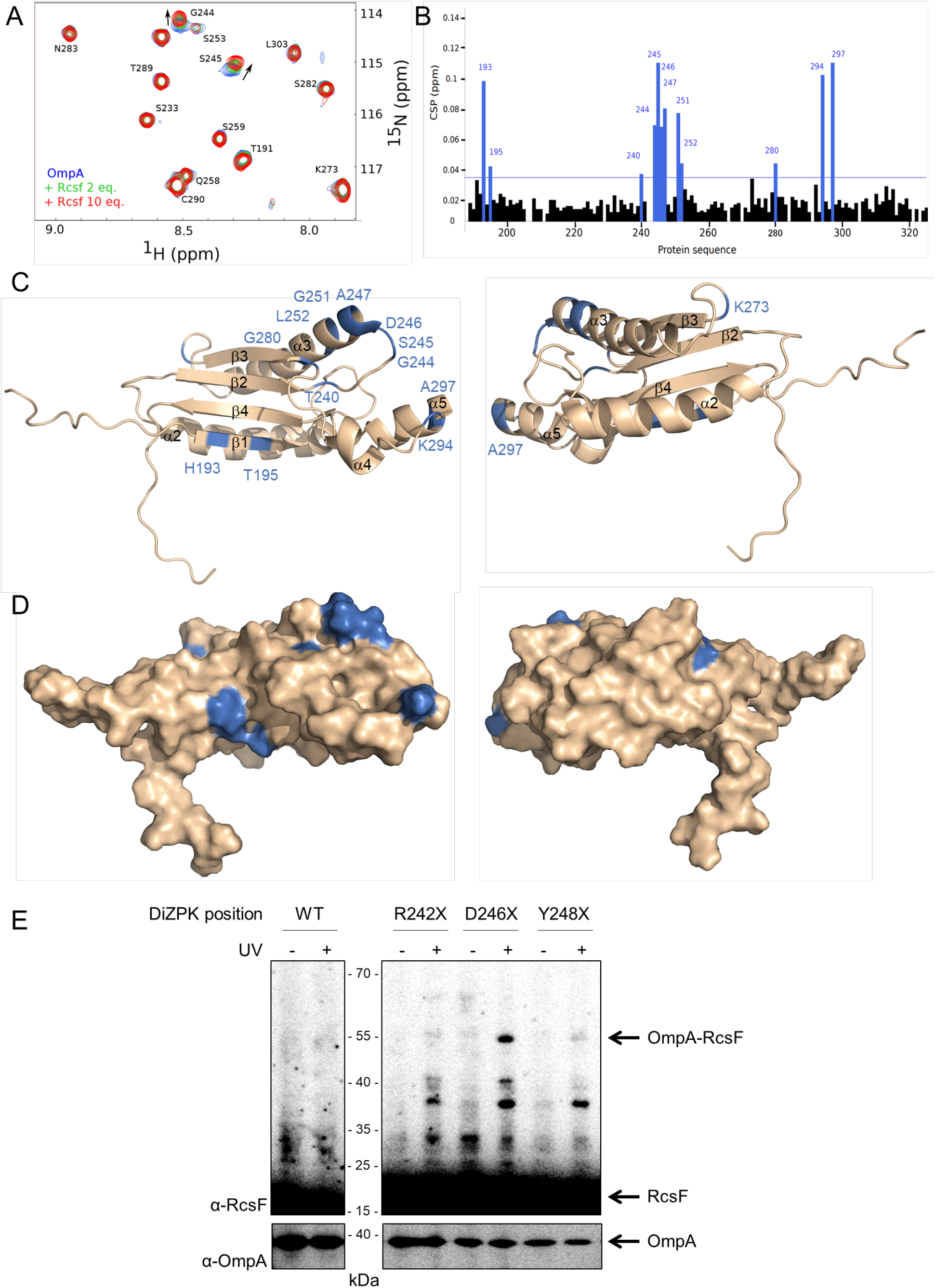
RcsF interacts with the periplasmic domain of OmpA *in vitro.* **(A)** An expanded region of the HSQC titration spectra of ^15^N-labeled OmpA_186-325_ with RcsF. Several residues in the ^1^H/^15^N BEST-TROSY-HSQC spectrum of OmpA (blue) show chemical shift perturbations (CSP) upon addition of RcsF at molar ratios of RcsF to OmpA of 2.5 (green) and 10 (red). Arrows indicate the direction of the chemical shift changes upon addition of RcsF to OmpA. **(B)** CSPs induced by the addition of RcsF to ^15^N-labeled OmpA_186-325_. Residues that showed CSP larger than two standard deviation are colored in blue. Cartoon **(C)** and surface **(D)** view of OmpA_186-325_ (PDB: 2MQE): top view (on the left) and bottom view (on the right). The residues in OmpA that undergo a CSP larger than two standard deviation (as in panel B) appear as blue and are labeled. Most of these residues are located between β-strand 2 and α-helix 3. **(E)** To confirm the importance of the loop between β-strand 2 and α-helix 3 of OmpA for the interaction with RcsF, we used site-specific photo-crosslinking. *ΔompA rcsF*^+^ cells expressing wild-type (WT) OmpA or three DiZPK-containing OmpA variants (OmpA_R242X_, OmpA_D246X_, and OmpA_Y248X_) from pPR21 were irradiated with UV light (+) or not (-), and protein samples were immunoblotted with an anti-RcsF antibody. A strong 55 kDa band, corresponding to the size of the OmpA-RcsF complex, was observed with the OmpA_D246X_ variant, confirming the NMR data. Weak complex formation was also observed with OmpA_R242X_ and OmpA_Y248X_. The expression levels of the OmpA variants were verified by immunoblotting (lower panel).

### IgaA and OmpA compete for RcsF across the periplasmic space

RcsF turns on the Rcs response by interacting with the periplasmic domain of the IM protein IgaA under stress (Hussein et al. 2018). Here, we found that the interaction between RcsF and OmpA takes place in the periplasm (Figure 5A), thus suggesting that OmpA and IgaA compete for binding RcsF across this compartment. To investigate this hypothesis, we determined the effect of artificially increasing the IgaA concentration on the formation of the IgaA-RcsF and OmpA-RcsF complexes. In these experiments, a triple Flag-tagged, functional version of IgaA (Hussein et al. 2018) was expressed from an inducible plasmid. To monitor complex formation, we carried out chemical crosslinking using bis(sulfosuccinimidyl)suberate (BS3), a bifunctional crosslinker. Remarkably, increasing the levels of IgaA led to changes in the IgaA-RcsF and OmpA-RcsF complexes that were inversely correlated: whereas overexpressing IgaA led to more IgaA-RcsF, it decreased the levels of the OmpA-RcsF complex (compare lanes 9 and 10 with lanes 7 and 8 in Figure 5B). Thus, IgaA and OmpA compete for RcsF across the periplasm. Because IgaA (200 copies per cell; (Li G-W 2014)) is far less abundant than OmpA (200,000 copies), even when overexpressed (we estimate that IgaA levels were increased 8-40 fold over baseline (Figure 5 – figure supplement 1; Materials and Methods) in the experiment above), these results suggested that IgaA has a substantially higher affinity for RcsF than OmpA. To probe this directly, we determined the affinity constants of the periplasmic domains of OmpA and IgaA for RcsF. Note that the periplasmic domain of IgaA interacts with RcsF *in vivo* and *in vitro* (Cho et al. 2014, Hussein et al. 2018)). First, from the NMR shift data, we calculated the equilibrium dissociation constant *(K_D_)* of OmpA_186-325_ for RcsF as being 125 ± 85 μM (Figure 5C). Second, using biolayer interferometry, we measured that the periplasmic domain of IgaA has a *K*_D_ of 1.6 ± 0.3 nM for RcsF (Figure 5D). Thus, these values confirm that IgaA has substantially more affinity for RcsF than OmpA (see Discussion). Interestingly, we noted that the BamA-RcsF complex was not modified by the increased expression of IgaA (lanes 7-10 in Figure 5B), which is consistent with the fact that BamA has a much higher affinity for RcsF (*K*_D_~400 nM; (Rodriguez-Alonso et al. 2020)) than OmpA.

**Figure 5.**
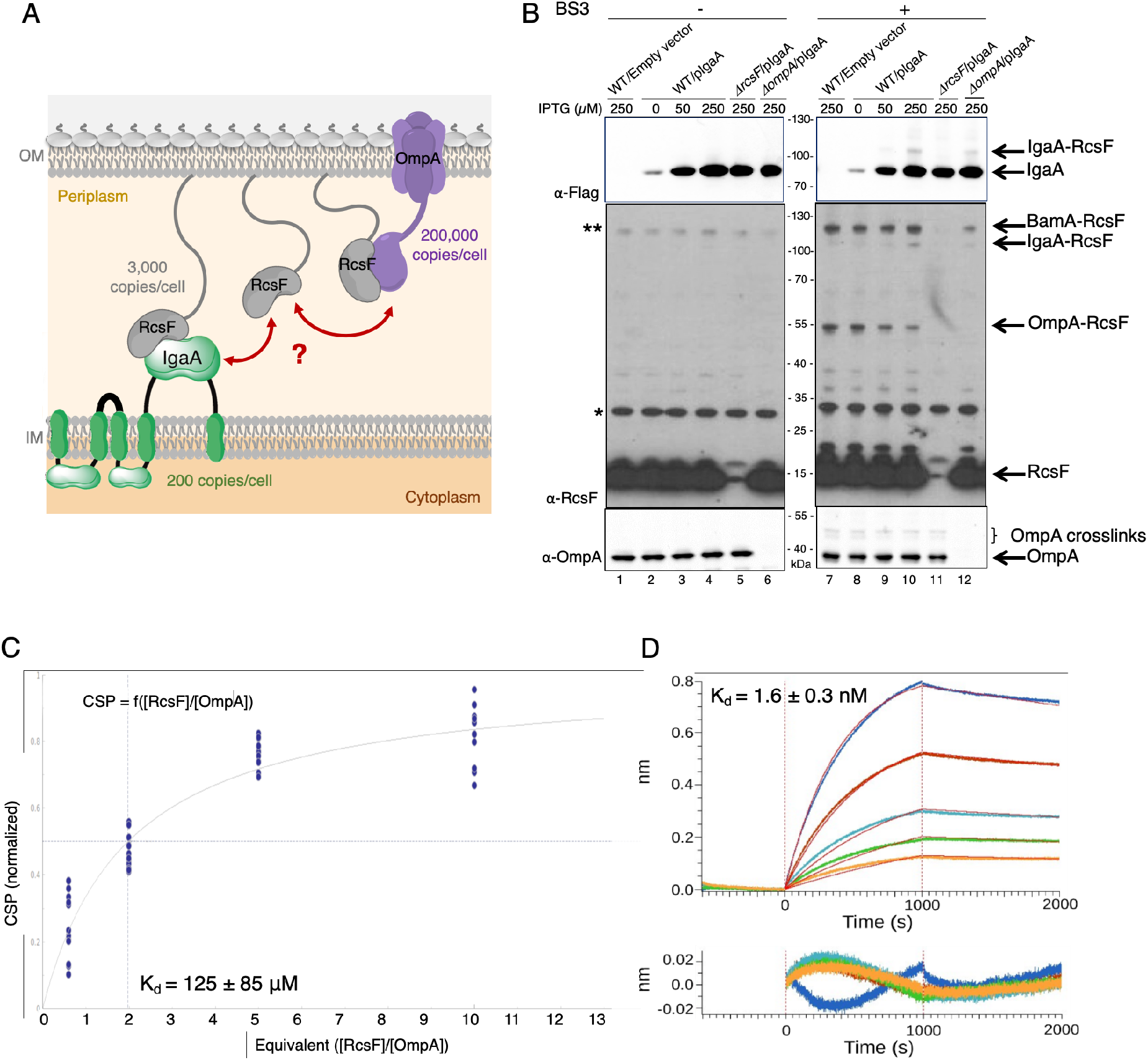
OmpA competes with the IM protein IgaA for RcsF. **(A)** Cartoon of the *E. coli* cell envelope, with IgaA in the IM and RcsF and OmpA in the OM. RcsF, an OM lipoprotein, interacts with OmpA. When in complex with RcsF, OmpA adopts its two-domain conformation, with an N-terminal β-barrel embedded in the OM and a C-terminal domain soluble in the periplasm. The periplasmic domain of OmpA interacts with RcsF, competing with IgaA for RcsF binding. Copy numbers are from (Li et al. 2014). **(B)** Impact of over-expressing IgaA on the OmpA-RcsF complex. Cells harboring pSC231, an empty vector used as control, or pIgaA (pSC237, expressing IgaA-Flag_3_ from an IPTG-inducible promoter) were harvested at mid-log phase and incubated without (lanes 1-6) or with (lanes 7-12) BS3. Protein samples were immunoblotted with α-Flag (upper panel), α-RcsF (middle panel), or α-OmpA_171-325_ (lower panel) antibodies. IgaA-Flag_3_ was expressed in WT (lanes 2-4 and 8-10), *ΔrcsF* (lanes 5 and 11), and *ΔompA* cells (lanes 6 and 12) with the indicated IPTG concentrations. * and **, non-specific bands detected by the polyclonal α-RcsF antibody. **(C)** Plot of the chemical shift perturbation (CSP) measured on OmpA resonances as a function of the RcsF:OmpA ratio. Only the residues with significant CSP (colored in blue in Fig. 4B) are plotted and used to fit the *K*_D_. **(D)** The interaction between RcsF and the periplasmic domain of IgaA was probed by biolayer interferometry (BLI). Sensortips carrying immobilized RcsF were dipped into increasing concentrations of IgaA (5.9, 8.9, 13.3, 20, 30 nM) from 0-1000 sec then into buffer (1000-2000 sec). Association and dissociation phases were fitted (red lines) to extract a *K_D_* value. Residuals from the fits are shown at the bottom of the panel.

## Discussion

### OmpA is unlikely the vehicle allowing RcsF to reach the surface

OmpA was first purified from *E. coli* membranes in 1977 (Chai and Foulds 1977) and has served as a model for OMP assembly since then. Although the predominant view is that OmpA folds into a two-domain conformation, with an N-terminal 8-stranded β-barrel and a C-terminal periplasmic domain, an alternative conformation in which OmpA forms a single, 16-stranded β-barrel, has been proposed to also exist (Singh et al. 2003, Stathopoulos 1996). Here, we integrated *in vivo* (Figures 2, 3, 4 and 5) and *in vitro* (Figures 4 and 5) approaches to dissect the interaction between OmpA and the lipoprotein RcsF; altogether, our data establish that OmpA is in the two-domain conformation in the OmpA-RcsF complex and that it is the C-terminal, periplasmic domain of OmpA that interacts with RcsF. Using protein-protein docking and molecular dynamics simulations, we built a three-dimensional model taking into accounts the results of the cross-linking experiments; in this model, RcsF and the periplasmic domain of OmpA show a remarkable surface complementarity (Figure 6). Interestingly, the peptidoglycan-binding region of OmpA (Park et al. 2012) remains accessible and is oriented in the direction opposite to the N-terminal end of RcsF. Our conclusions have two important implications.

**Figure 6.**
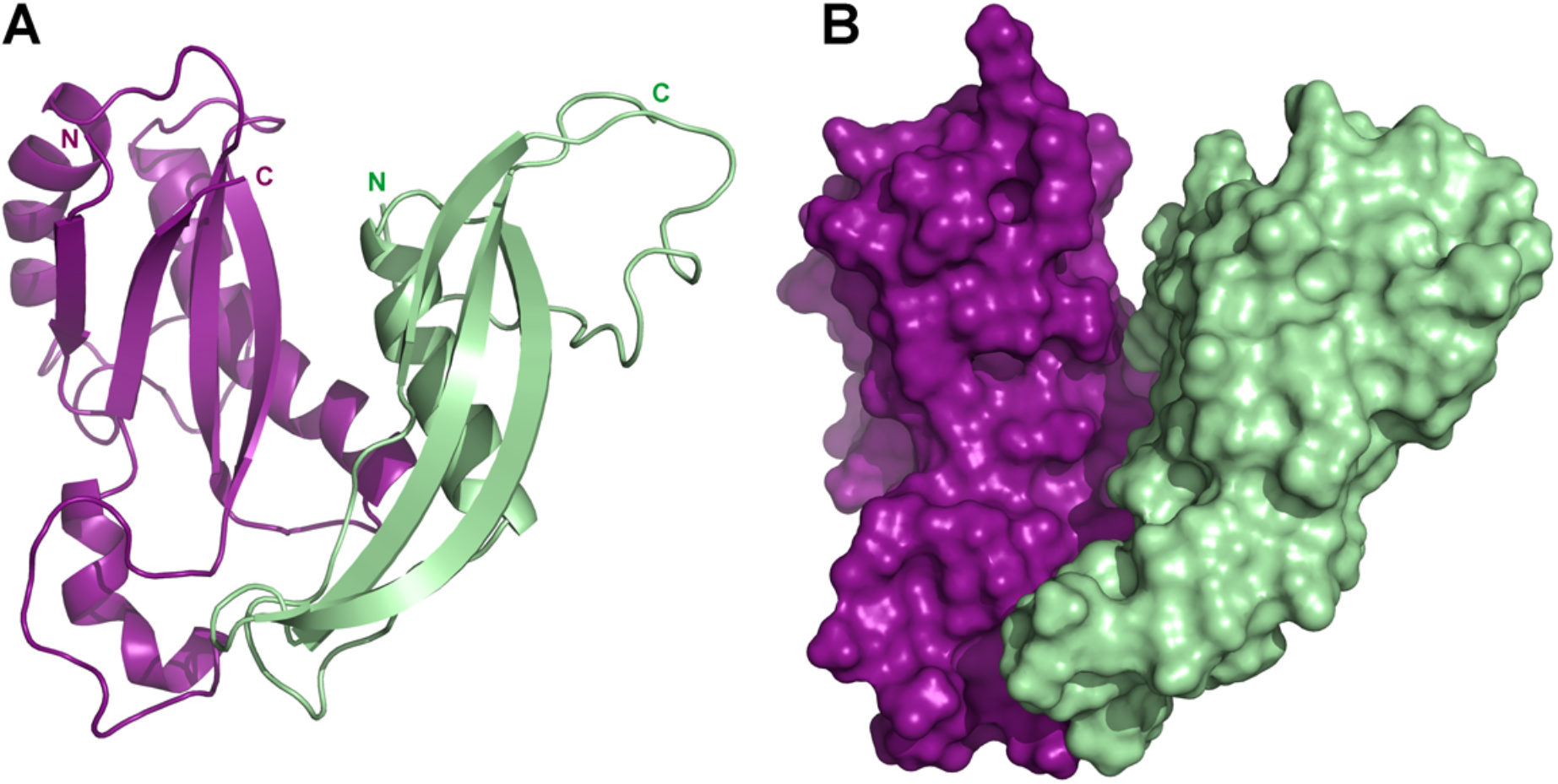
Model of the complex between RcsF and the C-terminal domain of OmpA. **(A)** Cartoon and (**B**) surface representation of a complex between RcsF (residues 44-134, colored in light green) and OmpA (residues 190-315, colored in purple). This model, which shows a remarkable surface complementarity between the two proteins, was obtained by protein-protein docking with constraints from the crosslinking experiments and further refined by all-atom molecular dynamics (50 ns). The N-terminal residue of RcsF points out in the same direction as the extremities of the periplasmic domain of OmpA, i.e. towards the OM, and this binding mode is compatible with the interaction of the periplasmic domain of OmpA with the peptidoglycan.

First, we and others previously showed that portions of RcsF are surface-exposed (Cho et al. 2014, Konovalova et al. 2014) and proposed OmpA, OmpC, and OmpF as vehicles for surface exposure (Cho et al. 2014, Konovalova et al. 2014). Understanding how RcsF reaches the surface is crucial: lipoprotein surface exposure is an emerging concept in *E. coli* and Enterobacteriaceae (Szewczyk and Collet 2016, Konovalova and Silhavy 2015) and how lipoproteins cross the OM remains to be clearly established. It was proposed that it is the N-terminal linker of RcsF that is exposed on the surface before being threaded through the lumen of OmpA and other OMPs (Konovalova et al. 2014). However, when OmpA adopts the two-domain conformation, as in the OmpA-RcsF complex, its β-barrel domain does not have an open channel (Figure 3 – figure supplement 1B) to accommodate the RcsF linker. We therefore conclude that RcsF does not reach the surface when in complex with OmpA, and predict that only OmpC and OmpF (which form large β-barrels) serve as vehicles for surface exposure. The fact that only a subset of the DiZPK-containing variants of RcsF that form a complex with OmpA also form a complex with OmpC/F (Figure 1) also supports the idea that the topology of OmpA-RcsF is different from that of OmpC/F-RcsF.

Second, a model was proposed in which RcsF, when in complex with OmpA, uses its positively charged, surface-exposed N-terminal linker to sense when interactions between lipopolysaccharide molecules are disturbed (Konovalova, Mitchell, and Silhavy 2016). However, if the RcsF linker is not surface-exposed in the OmpA-RcsF complex, as we show here, then the function of OmpA-RcsF in Rcs needs to be re-evaluated (see below). Importantly, we found that lipopolysaccharide alterations induce Rcs in MG1655 cells lacking *ompA* (Figure 6 – figure supplement 1), in contrast to what was previously reported in another strain background (MC4100) (Konovalova, Mitchell, and Silhavy 2016), which further questions the potential role of OmpA-RcsF in sensing lipopolysaccharide defects.

It will also be interesting to investigate the role of the BAM machinery in the assembly of OmpA-RcsF. It was reported that the formation of the OmpA-RcsF complex was likely mediated by BAM during the assembly of OmpA (Cho et al. 2014, Konovalova et al. 2014). However, our finding that RcsF interacts with the C-terminal, periplasmic domain of OmpA, whose folding is unlikely to depend on BAM (Noinaj, Gumbart, and Buchanan 2017), questions this conclusion. Further experiments, including pulse-chase experiments, will be carried out to clarify this point. Given that BAM both interacts with RcsF and assembles OmpA in the OM, we anticipate that interpreting the data will be challenging, and therefore these experiments are outside the scope of this publication.

### OmpA functions as a buffer for RcsF

Finally, if OmpA does not allow RcsF to reach the surface, what could be the function of the OmpA-RcsF complex? On the basis of our results, we propose that the role of OmpA in Rcs is to modulate the activity of this system. Consider the equilibrium dissociation constants of the IgaA-RcsF and OmpA-RcsF complexes:

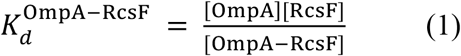

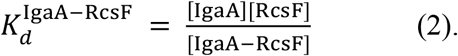

We estimate that OmpA’s C-terminal domain is present in the periplasm at ~1 mM, RcsF at ~15 μM, and IgaA at ~1 μM (Materials and Methods). If we take into account these estimates, then the following equation can be derived from (1) and (2) (Materials and Methods):

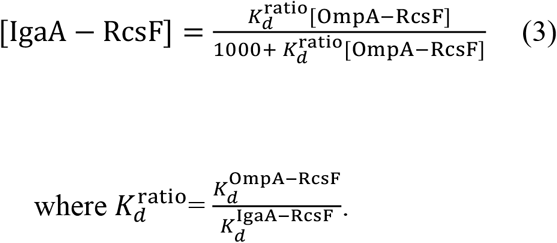

Because OmpA is highly abundant, [OmpA] can be considered constant. Thus, according to equation (1), the concentration of the OmpA-RcsF complex increases linearly with the concentration of RcsF in the periplasm (Figure 6 – figure supplement 2). From equation (3), we conclude that the concentration of IgaA-RcsF will also increase as a function of [RcsF] in the periplasm. However, in this case, the increase is not linear, but is decelerated and follows a hyperbolic curve (Figure 6 – figure supplement 2), indicating that OmpA functions as a buffer for RcsF, negatively impacting its ability to activate Rcs. The ~80,000-fold difference in affinity for RcsF that we measured between IgaA and OmpA (Figure 5) does not seem to support this conclusion: with such a high affinity for RcsF, IgaA should always outcompete OmpA and formation of the IgaA-RcsF complex should be constitutive, which is not the case (Hussein et al. 2018). To explain the apparent discrepancy between the *in vivo* observations and the *in vitro* measurements, we propose that intracellular factors such as attachment of the periplasmic domains of these proteins to their respective membrane anchors (we used the soluble periplasmic domains of OmpA, IgaA and RcsF for the *in vitro* measurements) and binding of OmpA to the peptidoglycan (Reusch 2012) modulate the affinity of OmpA and IgaA for RcsF *in vivo*, allowing them to actually compete for RcsF. It is remarkable that OmpA, a protein that had mostly been known for its role in stabilizing the OM, is also involved in the network controlling the activity of Rcs, one of the most complex signal transduction systems in bacteria (Cho et al. 2014, Hussein et al. 2018, Laloux and Collet 2017), whose correct functioning is critical for success of commensal Enterobacteriaceae and virulence of pathogens.

## Materials and Methods

### Bacterial strains, primers, and plasmids

The bacterial strains and primers used in this study are listed in Supplementary Tables 1 and 2, respectively. The parental *E. coli* strain DH300 is a MG1655 derivative containing a deletion of the *lac* region; it also carries a chromosomal *rprA::lacZ* fusion at the λ phage attachment site to monitor Rcs activation (Majdalani, Hernandez, and Gottesman 2002). The *ompA, ompR* and *rcsF* deletion mutants were obtained by transferring the corresponding alleles from the Keio collection (kan^R^) (Baba et al. 2006) into DH300 (Majdalani, Hernandez, and Gottesman 2002) *via* P1 phage transduction, which was verified *via* PCR. To excise the kanamycin-resistance cassette, we used pCP20 as previously described (Datsenko and Wanner 2000). To insert *ompA-His* (encoding OmpA with six histidines at the C-terminus), *ompA_TH189_-His, ompA_TH243_-His,* and *ompX-ompA_171-325_* on the chromosome at the *ompA* locus, we performed λ-Red recombineering (Yu et al. 2000) with the pSIM5-Tet plasmid (Koskiniemi et al. 2011). First, a *cat-sacB* cassette encoding chloramphenicol acetyl transferase (cat) and SacB, a protein conferring sensitivity to sucrose, was amplified from strain CH1990 using primers “ompA_delCmSB F” and “ompA_delCmSB R”. The resulting PCR product shared 40 bp of homology to the 5’ UTR and 3’ UTR of *ompA* at its 5’ and 3’ ends, respectively, and was used for λ-Red recombineering (Yu et al. 2000). We selected transformants for chloramphenicol resistance and verified that the *cat-sacB* cassette replaced *ompA* by sequencing across the junctions. The *cat-sacB* cassette was subsequently replaced by one of the versions of *ompA* (*ompA-His, ompA_TH189_-His, ompA_TH243_-His*, or *ompX-ompA_171-325_*) using λ-Red recombineering and negative selection on sucrose-containing medium (Thomason et al. 2014, Gennaris et al. 2015). *ompA-His, ompA_TH189_-His,* and *ompA_TH243_-His* were amplified *via* PCR using primers “cmSB to OmpA_F” and “cmSB to OmpAhis_R” and plasmids pPR11, pPR11_189thrombin_, and pPR11_243thrombin_ (see below) as templates, respectively. The PCR product of *ompX-OmpA_171-325_* was generated using primers “JLEo20-F-Chrom-OmpX” and “JLEo22-R-Chrom-OmpACt”, and pJLE17-OmpX-OmpACter as template. Strains were verified through DNA sequencing. To delete *ompA_171-325_,* we performed λ-Red recombineering on the chromosome at the *ompA* locus in a way similar to the description above. Briefly, the kanamycin cassette from the strain SEN588 *(ΔompA::kan)* was PCR-amplified using primers “ompAc_delKm F” and “ompAc_delKm R” to encompass the flanking regions of *ompA_171-325_*. We selected transformants for kanamycin resistance.

The plasmids used in this study are described in Supplementary Table 3. pSC253, encoding RcsF, was constructed *via* digestion of pSC202 (Cho et al. 2014) with *KpnI* and *HindIII* and insertion of the generated fragment into pBAD18 (Guzman et al. 1995). To generate the *rcsF* variants containing an amber codon (TAG) at selected positions, site-directed mutagenesis was performed on pSC253 using primers described in Supplementary Table 2. To generate pPR11, *ompA-His* was PCR-amplified using primers “OmpA(NcoI) F” and “OmpA-his(XbaI) R” and chromosomal *E. coli* DNA as template. The PCR product was inserted into pDSW204 restricted with, *Nco*I and *Xba*I, generating pPR11. To insert thrombin cleavage sites after residues 189 and 243 in OmpA, site-directed mutagenesis was performed using primer pair “189VPRGS thr_F” and “189VPRGS thr_R” and primer pair “243LVPR thr_F” and “243LVPR thr_R”, respectively, on pPR11, yielding pPR11_189thrombin_ [OmpA(189-Val-*Val-Pro-Arg-Gly-Ser*-Gln-190)] and pPR11_243thrombin_ [OmpA(243-Ile-*Leu-Val-Pro-Arg*-Gly-244)], respectively. pJLE17-OmpX-OmpA_171-325_ (encoding OmpAX) was constructed as follows. The two PCR products corresponding to *ompX* and *OmpA_171-325_* were generated using primer pair “OmpX(NcoI)F” and “OmpX-OmpAc_R” and primer pair “OmpX-OmpAc_F” “JLEo16-R-XbaI-OmpACt”, respectively, and chromosomal *E. coli* DNA as template. To join the two PCR fragments, overlapping PCR was performed using primers “OmpX(NcoI)F” and “JLEo16-R-XbaI-OmpACt”, generating the PCR product encoding OmpA_X_. Next, the PCR product of *OmpA_X_* was digested with *Nco*I and *Xba*I and ligated with pDSW204 pre-digested with *Nco*I and *Xba*I, yielding pJLE17-OmpX-OmpACter. pKiD22, expressing OmpA_Pal_, was constructed as follows. The sequence encoding the signal sequence and lipobox of the lipoprotein Pal was PCR-amplified using primers “Palss(NcoI)F” and “Palss-OmpAc_R” and the sequence encoding OmpA_171-325_ using primers “Palss-OmpAc_F” and “OmpA_stop_Flag_3_(KpnI)R”. Chromosomal *E. coli* DNA was used as template. To join the two PCR fragments, overlapping PCR was performed using primers “Palss(NcoI)F” and “OmpA_stop_Flag_3_(KpnI)R”. The PCR product encoding ssPal-OmpA_171-325_ was digested with *Nco*I and *Kpn*I and ligated with pDSW204, pre-digested with *Nco*I and *Kpn*I, yielding pKiD22.

To generate pPR21, site-directed mutagenesis was performed on pPR11 using primers “OA_stopTGA_F” and “OA_stopTGA_R” to insert a stop codon upstream of the 6xHis tag. To generate the *ompA* variants containing an amber codon (TAG) at selected positions, site directed mutagenesis was performed on pPR21 using primers described in Supplementary Table 2. To obtain pKiD5, OmpA_186-325_ with an N-terminal Strep-tag but no signal sequence was PCR-amplified using primers “KiDo14-F-NdeI-Strep-OmpACTD” and “KiDo15-R-SacI-Rbs-OmpACTD” and chromosomal *E. coli* DNA as template. The PCR product was digested with *Nde*I and *Sac*I and inserted into pET21a. To prepare a version of pAM238 containing *lac1*^q^, a *trc* promoter, and a triple Flag tag (Flag_3_), we prepared a PCR product using primers “lacIq NsiI_F” and “Flag3-KpnI_R” and pMER77 as template (Hemmis et al. 2011). This product was digested with *Nsi*I and *Kpn*I and ligated with pAM238 pre-digested with *Pst*I and *Kpn*I. To reduce the basal activity of the *trc* promoter, we modified the −10 region (from TATAAT to CATTAT) and the −35 region (from TTTACA to TTGACA), generating pSC231 (Weiss et al. 1999). The coding sequences for OmpA and IgaA were obtained by PCR-amplification using primer pair “OmpA(NcoI) F” and “OmpA XbaI flag R” and primer pair “IgaA(NcoI) F” and “IgaA XbaI Flag3 R”, respectively. Each product was digested with *Nco*I and *Xba*I and ligated into pSC231 pre-digested with the same enzymes, generating pPR4 and pSC237, respectively.

### *In vivo* site-specific photo-crosslinking using DiZPK

We used the site-specific photo-crosslinking method described previously (Cho et al. 2014) with some modifications. To incorporate *N*^6^-((3-(3-methyl-3*H*-diazirin-3-yl)propyl)carbamoyl)-*L*-lysine (DiZPK) into RcsF, we used the pSup-Mb-DIZPK-RS plasmid encoding an evolved *Methanosarcina barkeri* pyrrolysyl-tRNA synthetase (PylRS) and an optimized 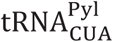 suppressor (Zhang et al. 2011). DH300 *DrcsF* (PL358) cells were cotransformed with pSup-Mb-DIZPK-RS and one of the plasmids containing an amber codon in *rcsF* in pSC253. Cells were grown in 3-(N-morpholino)propanesulfonic acid (MOPS) minimal medium supplemented with 0.2% glucose, 0.2% L-arabinose (MOPS-glucose/arabinose minimal medium), and 0.8 mM DiZPK (no supplement of other amino acids; see the reasons for using MOPS medium below) (Neidhardt, Bloch, and Smith 1974). Cell cultures were grown to an OD_600_ of 1-1.2 and 1 mL of samples was irradiated with UV light at 365 nm or left unirradiated for 10 min. Cells were precipitated with trichloroacetic acid (TCA), and the pellets were washed with acetone and solubilized in 60 μL of SDS-sample buffer before further analysis. We used a similar method to incorporate DiZPK into OmpA with minor modifications; DH300 *DompA* (PR46) cells were co-transformed with pSup-Mb-DIZPK-RS and one of the plasmids containing an amber codon in *ompA* in pPR21. Cells were grown in LB medium supplemented with 0.2% L-arabinose, 200 μM IPTG and 1 mM DiZPK. Cell cultures were grown to an OD_600_ of 1 and 0.75 mL of samples was irradiated with UV light at 365 nm or left unirradiated for 10 min. Cells were precipitated with TCA, and the pellets were washed with acetone and solubilized in 100 μL of SDS-sample buffer before further analysis.

In previous experiments incorporating *p*-benzoyl-L-phenylalanine into RcsF, we used LB as the growth medium (Cho et al. 2014). However, we found that the expression levels of the DiZPK-containing RcsF mutant proteins were substantially lower when cells were grown in LB (data not shown); in contrast, the expression levels of RcsF were greatly enhanced in MOPS-glucose/arabinose minimal medium (data not shown). Therefore, we used MOPS-glucose/arabinose minimal medium for all photo-crosslinking experiments involving DiZPK-containing RcsF variants.

### Synthesis of DiZPK

DiZPK was synthesized as described previously (Zhang et al. 2011).

### Immunoprecipitation of RcsF-containing complexes and thrombin cleavage

Protein samples in SDS-sample buffer (without reducing agent) were denatured for 15 min at 65 °C and 15 min at 95 °C with vigorous shaking. Next, the samples were diluted in 750 μL of KI buffer (50 mM Tris-HCl [pH 8], 2% Triton X-100, 150 mM NaCl, 1 mM EDTA) and centrifuged at 4 °C for 3 min at 12,000 x *g* to harvest the PG. Photo-crosslinked RcsF complexes were immunoprecipitated by adding 1 μL of undiluted guinea pig anti-RcsF antibody (Cho et al. 2014) and 10 μL of protein A/G magnetic beads (Pierce™); samples were incubated for 1 h on a wheel at room temperature. After three washes with 500 μL of KI buffer, RcsF complexes were eluted with 20 μL of glycine buffer (100 mM glycine [pH 1.5], 0.1% Triton X-100) after 10 min of incubation at room temperature. Proteins samples were neutralized with 2 μL of 1.5 M Tris-HCl [pH 8.8] and diluted with 18 μL of KI buffer before SDS-PAGE and immunoblotting or thrombin cleavage. For thrombin cleavage, 20 μL of the elution samples were incubated for 1 h at room temperature with 1 μL of thrombin (thrombin from bovine plasma, Sigma). Samples were analyzed *via* SDS-PAGE and immunoblotting as indicated in the figure legends.

### Immunoblotting and antibodies

Protein bands were transferred from the gels onto nitrocellulose membranes (Millipore) using a semi-dry electroblotting system. The membranes were blocked with 5% skim milk. The rest of the immunoblot steps were performed using standard protocols. Signal from antibody binding was visualized by detecting chemiluminescence from the reaction of horseradish peroxidase with luminol. Polyclonal RcsF antibodies were purified against the carboxy-terminal domain of RcsF as previously described (Cho et al. 2014) and used at a dilution of 1:20,000 in 1% skim milk in 50 mM Tris-HCl [pH 7.6], 0.15 M NaCl and 0.1% Tween 20 (TBST). Since we found that the anti-OmpA antibody only recognizes the periplasmic domain (data not shown), we used an antibody directed against loop 4 of the β-barrel of OmpA to detect OmpA_1-170_. The anti-OmpA antibodies are gifts from the Lloubes and Bernstein laboratories (Cascales et al. 2002, Hussain and Bernstein 2018). These antibodies were used at dilutions of 1:20,000 and 1:10,000, respectively. The anti-His antibody conjugated to horseradish peroxidase (Qiagen) was used at a dilution of 1:5,000. The anti-Flag M2 monoclonal antibody (F1804, Sigma) was used at a dilution of 1:20,000.

### *In vivo* 3,3’-dithio-bis[sulfosuccinimidylpropionate] and bis(sulfosuccinimidyl)-suberate (BS3) crosslinking

DTSSP and BS3 (CovaChem) are bifunctional primary amine crosslinkers; DTSSP contains a disulfide bond in its spacer arm. *In vivo* crosslinking has been performed as described previously (Cho et al. 2014). The media used are MOPS-glucose minimal medium (Neidhardt, Bloch, and Smith 1974) (Figure 3) and LB (Figure 5)

### β-galactosidase assay

*E. coli* cells were grown in MOPS-glucose minimal medium (Neidhardt, Bloch, and Smith 1974) or LB to mid-log phase (OD_600_ = 0.4-0.6). β-galactosidase activity was measured as described previously (Zhang and Bremer 1995).

### Expression and purification of RcsF, OmpA_186-325_ and of the periplasmic domain of IgaA

Expression and purification of RcsF with a C-terminal His-tag were performed as previously described (Leverrier et al. 2011). *E. coli* BL21(DE3) cells harboring pKiD5 expressing N-terminal Strep-tagged OmpA_186-325_ were grown at 37°C in M9-glucose minimal medium containing 1 g/L ^15^NH_4_Cl (99%, ^15^N; Eurisotop) to uniformly label the protein with ^15^N. The expression was induced by adding 1 mM IPTG at an OD_600_ of 0.5. After a 5h induction, cells were harvested by centrifugation and resuspended in 20 mL of lysis buffer (20 mM Tris-HCl, 500 mM NaCl [pH8], containing a protease inhibitor cocktail (Complete^®^, Roche)). Resuspended cells were stored at −20 °C. Frozen cells were thawed on ice and lysed by two passages through a French pressure cell at 1,500 psi. The soluble fraction was isolated after centrifugation for 1h at 40,000 x *g* at 4 °C. The supernatant was filtered through 0.45 μm filters and loaded onto a 5 mL Strep-Tactin column (IBA, Lifesciences), previously equilibrated in buffer A (50 mM NaH_2_PO_4_, 100 mM NaCl [pH7.0]). After a washing step with buffer A, elution was performed with buffer A supplemented with 2.5 mM D-desthiobiotin. The sample was then further purified using size-exclusion chromatography on a HiLoad 16/60 Superdex 75 column (GE Healthcare) using buffer A. To express IgaA_361-654_ with a C-terminal His-tag, *E. coli* SHuffle T7 cells harboring pSC211 were grown at 37°C in LB. Expression of the protein was induced by adding 1 mM IPTG at an OD_600_ of 0.5. After a 5h induction, cells were harvested by centrifugation and resuspended in 20 mL of lysis buffer. Re-suspended cells were stored at −20 °C. Frozen cells were thawed on ice and processed as explained above for Strep-tagged OmpA_186-325_. IgaA_361-654_ was purified using Ni-NTA agarose beads (5 mL; IBA Lifescience), previously equilibrated in buffer B (20 mM NaH_2_PO_4_, 150 mM NaCl [pH7.5]). After washing the resin with buffer B supplemented with 20 mM imidazole, proteins were eluted with five column volumes of buffer B supplemented with 200 mM imidazole. As a final purification step, a size-exclusion chromatography was performed using a HiLoad 16/60 Superdex 200 column (GE Healthcare) with buffer B Purity was checked via SDS-PAGE with Coomassie Staining and concentration was performed using Vivaspin Turbo apparatus (Sartorius) with a 5 kDa molecular weight cut-off.

### NMR titration experiments of ^15^N-labelled OmpA_186-325_ with RcsF

50 μM ^15^N-labelled OmpA_186-325_ in buffer A at 25°C was titrated with increasing concentrations of unlabelled RcsF in buffer A to reach 2 and 10 RcsF/OmpA_186-325_ molar ratios. To follow ^15^N-^1^H resonances chemical shifts perturbations, ^15^N-^1^H BEST-TROSY-HSQC correlation experiments were recorded at 25°C using Bruker AVANCE spectrometer operating at 700 MHz proton frequency equipped with a TCI cryoprobe (Favier and Brutscher 2011). OmpA_186-325_ assignments were transposed from BMRB entry 25030. Chemical shift perturbations (CSP) were calculated on a per-residue basis for the highest substrate-to-protein ratio as described previously (Egan et al. 2018). Spectra were processed with Topspin 3.57 (Bruker) and analyzed with ccpnmr 3 (https://www.ccpn.ac.uk).

### Biolayer interferometry

Biolayer Interferometry Experiments (BLI) were recorded on an OctetRED96e (Fortebio) using streptavidin (SA) biosensors (Fortebio). To biotinylate RcsF, RcsF-His (5mg/mL in 0.1 M MES [pH 5.5]) was incubated with Biotin-LC-Hydrazide (1.25 mM, final concentration; Sigma) and N-(3-Dimethylaminopropyl)-N’-ethylcarbodiimide (EDC; 6.5 mM, final concentration) during 2h at 22°C under agitation. Biotininylated RcsF was dialysed against buffer C (10 mM Hepes, 150 mM NaCl [pH7.5]) and immobilized at 5 μg/mL onto SA tips in buffer C supplemented with 0.02% Tween-20 to reach ~3.5 nM of immobilization level. RcsF-loaded biosensors were dipped into different concentrations of the periplasmic domain of IgaA (5.9 nM to 30 nM) in buffer C at 23°C. Kinetics were recorded with 1000 sec association and 1000 sec dissociation phases, and repeated 4 times with 10mM HCl pulses (18 sec in total) used for regeneration between cycles. Sensorgrams were subtracted for contribution of buffer alone and binding of non-functionalized biosensors. Kinetic analysis of the data was performed using 1:1 interaction model in the ForteBio data analysis software. *K_D_* obtained from 4 independent injection series were averaged and produced a *K_D_* of 1.6 nM with a standard deviation of 0.3.

### Estimation of the expression levels of IgaA

Overexpressed IgaA has a triple Flag tag (IgaA-Flag_3_). To compare the expression levels of IgaA to those of OmpA, we generated a triple Flag-tagged version of OmpA (OmpA-Flag_3_); this variant was expressed from an IPTG-inducible plasmid in the *ompA* strain. If we compare the intensity of the signal corresponding to OmpA-Flag_3_ detected either by the anti-OmpA or the anti-Flag antibodies, we estimate the anti-Flag antibodies to be ~25 times more sensitive than the anti-OmpA antibodies (lanes 1-4 in Figure 5 – figure supplement 1). With this value in hand, we can now compare the expression levels of IgaA-Flag_3_ to those of untagged OmpA produced from the chromosome and detected with the anti-OmpA antibodies (lanes 5-9 in Figure 5 – figure supplement 1); because the intensities of OmpA and IgaA-Flag_3_ in lane 8 are similar, we estimate that IgaA-Flag_3_ is 25 times less abundant than OmpA when IgaA is expressed with 250 μM IPTG; likewise, we estimate the levels of IgaA-Flag_3_ to be ~1/125 (0.8%) of those of OmpA when IgaA is expressed with 50 μM IPTG. Because the concentration of OmpA is ~1 mM, we estimate the concentration of overexpressed IgaA to be 40 and 8 μM, respectively.

### Molecular modelling

HADDOCK 2.4 web server (van Zundert et al. 2016, Wassenaar et al. 2012) was used for protein-protein docking using the structures 2MQE for OmpA-CTD (Ishida, Garcia-Herrero, and Vogel 2014) and 2L8Y for RcsF (Rogov et al. 2011). Six residues identified from the crosslinking experiments were considered as active during the docking calculations: 242, 246 and 248 for OmpA and 45, 79 and 89 for RcsF. The resulting clusters were inspected visually, and the one compatible with the interaction between the periplasmic domain of OmpA and the peptidoglycan (Park et al. 2012) was selected for further refinement using molecular dynamics. Molecular dynamics simulations were carried out with GROMACS version 2020.1 (Abraham et al. 2015) using the OPLS-AA (Jorgensen, Maxwell, and Tirado-Rives 1996) force field. Each system was energy-minimized until convergence using a steepest descents algorithm. Molecular dynamics with position restraints was then performed (50 ps NVT and 50 ps NPT), followed by the production run of 50 ns. During the position restraints and production runs, the V-rescale and Parrinello-Rahman methods were used for temperature and pressure coupling, respectively. Electrostatics were calculated with the particle mesh Ewald method. The P-LINCS algorithm was used to constrain bond lengths, and a time step of 2 fs was used throughout.

### Estimation of the concentrations of OmpA, RcsF and IgaA in the periplasm

According to (Li et al. 2014), each cell contains ~200,000 molecules of OmpA in rich media, which corresponds to ~3.448 x 10^-19^ mol (200,000/Avogadro constant). If we consider the volume of an *E. coli* cell to be 10^-18^ m^3^ (10^-15^ L) and compare it to a cube with 1 μm edges (*E. coli* has 0.5 μm in width and 2 μm in length; EcoliWiki ecoliwiki.net/colipedia/index.php/Escherichia_coli), we calculate that the cellular concentration of OmpA is ~3.448 x 10 ^-4^ M. The volume of the envelope being ~30% of the total cell volume (Stock, Rauch, and Roseman 1977), we calculate that the concentration of the C-terminal domain of OmpA in the periplasm is ~1 mM. If we perform the same calculations for RcsF and IgaA (~3,000 and ~200 copies/cell, respectively), we find that their periplasmic concentrations are ~15 μM and ~1 μM, respectively.

### OmpA functions as a buffer for RcsF

The equilibrium dissociation constants for the OmpA-RcsF and IgaA-RcsF complexes are:

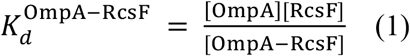

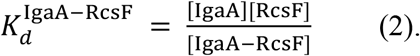

If we divide (1) with (2), we obtain:

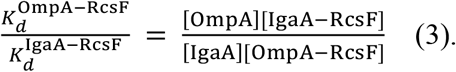

If we replace 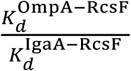 by 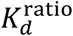, then we can rearrange equation (3) to

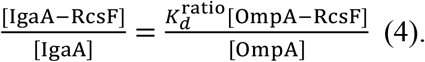

Under physiological conditions, we estimate the concentration of IgaA to be ~1 μM (see above). Thus, because [IgaA] + [IgaA-RcsF] = 1 μM, equation 4 can be successively rearranged to

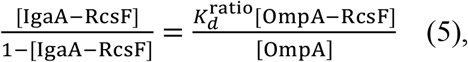

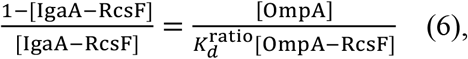

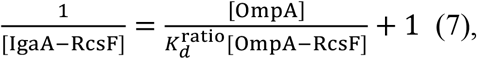

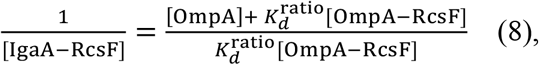

yielding:

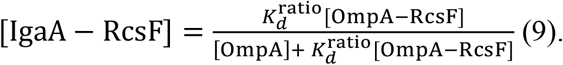

In equation (1), [OmpA], which is ~1000 μM, can be considered as a constant. Therefore, equations (1) and (9) become:

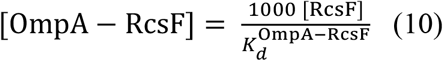

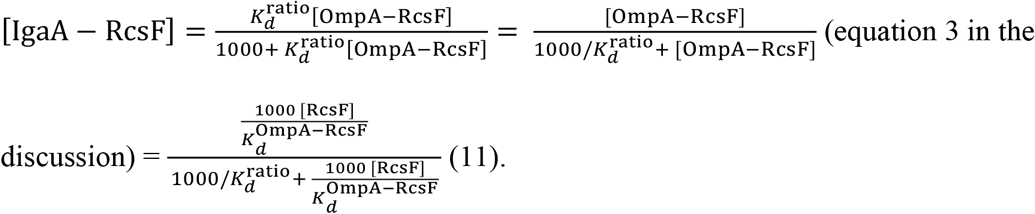

Thus, from equation (11), we conclude that whereas [OmpA-RcsF] increases linearly to [RcsF], [IgaA-RcsF] increases proportionally, but not linearly, to [RcsF]. Thus, OmpA functions as a buffer for RcsF.

### Analysis of protein structures

Protein structures were downloaded from the Protein Data Bank (http://www.rcsb.org; PDB codes are indicated) and visualized using PyMOL Molecular Graphics System (Version 2.3.4, Schrödinger, LLC). FASTA protein sequences were downloaded from Uniprot (http://www.uniprot.org/).

## Acknowledgments

We are indebted to Marie Renault (CNRS, Toulouse) for discussion and to Harris Bernstein (NIH, USA), Eric Cascales and Roland Lloubes (CNRS, Marseille) for sharing antibody with us. We are grateful to Alexandra Gennaris (UCLouvain), Pauline Leverrier (UClouvain) and Camille Goemans (EMBL, Heidelberg) for critically reading the manuscript and providing feedback. We thank the members of the lab for helpful discussions, and A. Boujtat for assistance in experiments. This work used the platforms of the Grenoble Instruct-ERIC center (ISBG; UMS 3518 CNRS-CEA-UGA-EMBL) within the Grenoble Partnership for Structural Biology (PSB), supported by FRISBI (ANR-10-INBS-05-02) and GRAL, financed within the University Grenoble Alpes graduate school (Ecoles Universitaires de Recherche) CBH-EUR-GS (ANR-17-EURE-0003). Authors acknowledge the SPR/BLI platform personal, Jean-Baptiste Reiser and Anne Chouquet, for their help and assistance. K.D. and R.B. are FRIA research fellows and J.L is “Chargée de Recherches” of the Fonds de la Recherche Scientifique FRS-FNRS. This work was supported by grants from FRFS-WELBIO, from the FRS-FNRS, and from the Fédération Wallonie-Bruxelles (ARC 17/22-087).

---

**Figure 1 — figure supplement 1.**
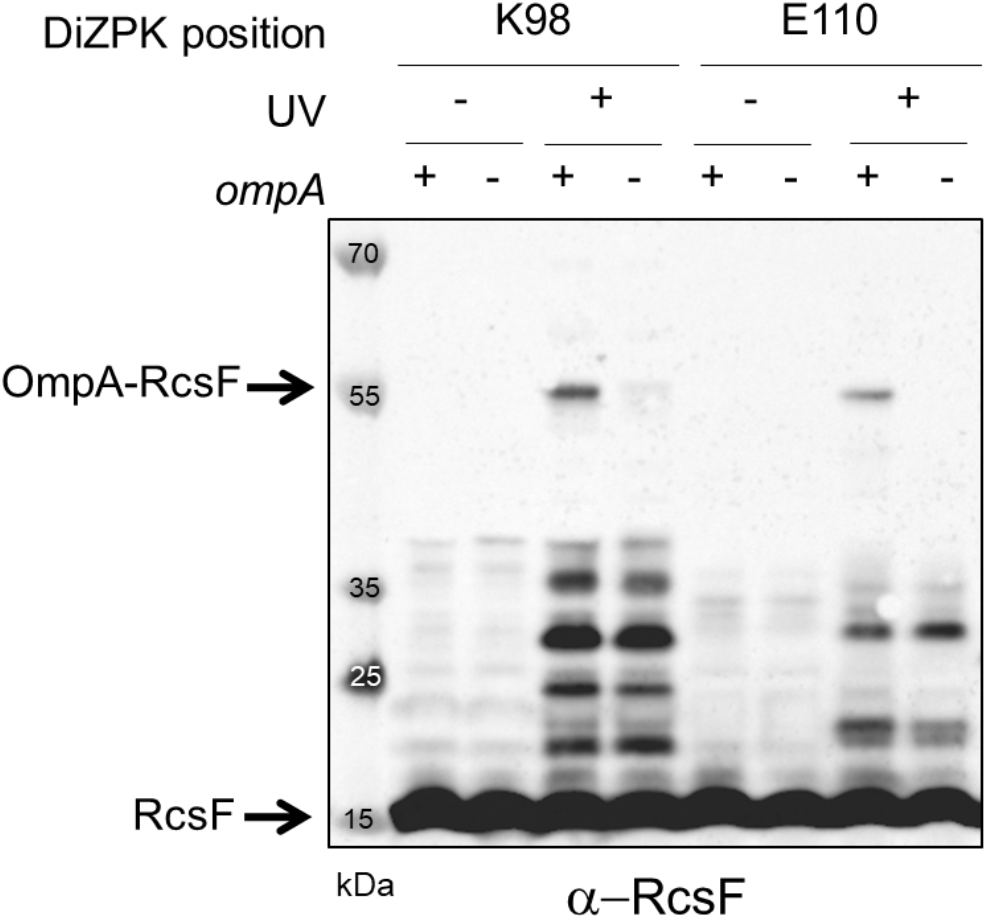
The UV-dependent 55 kDa band is the OmpA-RcsF complex. To confirm the identification of the 55 kDa band as OmpA-RcsF, two DiZPK-containing RcsF variants (K98 and E110) were expressed from pSC253 in *ΔrcsF* and *ΔrcsFΔompA* cells. Following UV exposure, the 55 kDa band only formed in cells in which OmpA was produced. Protein samples were immunoblotted with an anti-RcsF antibody.

**Figure 1 — figure supplement 2.**
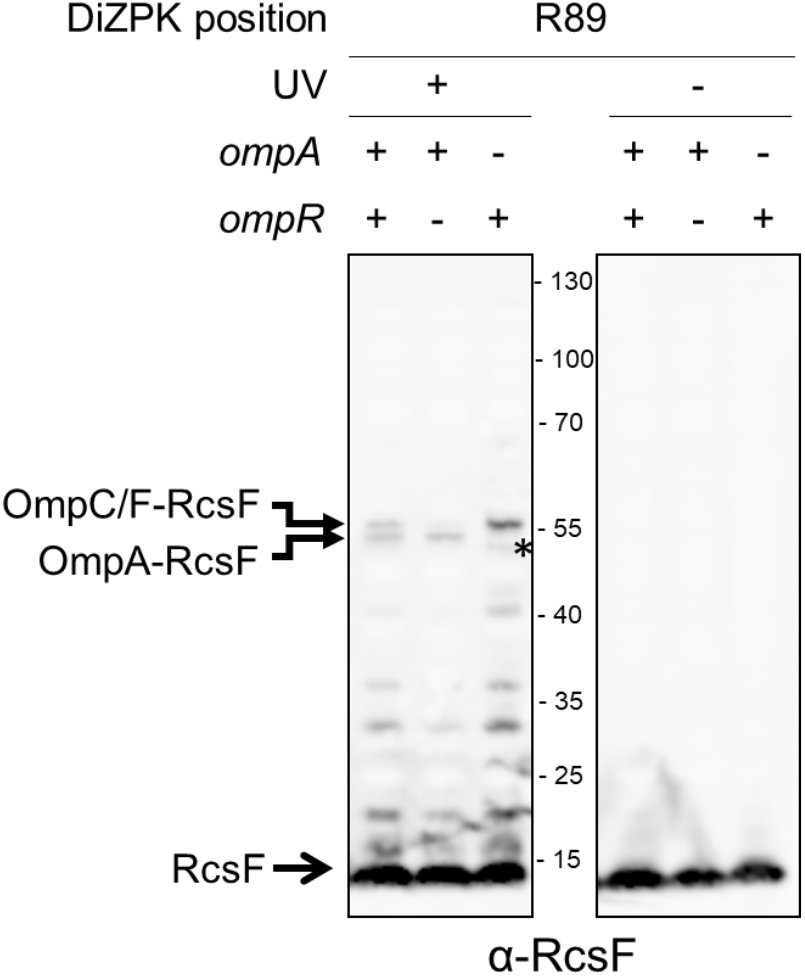
The band migrating above the OmpA-RcsF complex is the OmpC/F-RcsF complex. A band migrating slightly higher than OmpA-RcsF was observed with the RcsF_R89X_ variant expressed from pSC253 in *ΔrcsF* cells following UV exposure. This band corresponds to the OmpC/F-RcsF complex because it did not form in cells lacking *ompR,* a transcription factor required for the production of OmpC and OmpF. The unknown band marked by * migrates lower than OmpA-RcsF.

**Figure 3 — figure supplement 1.**
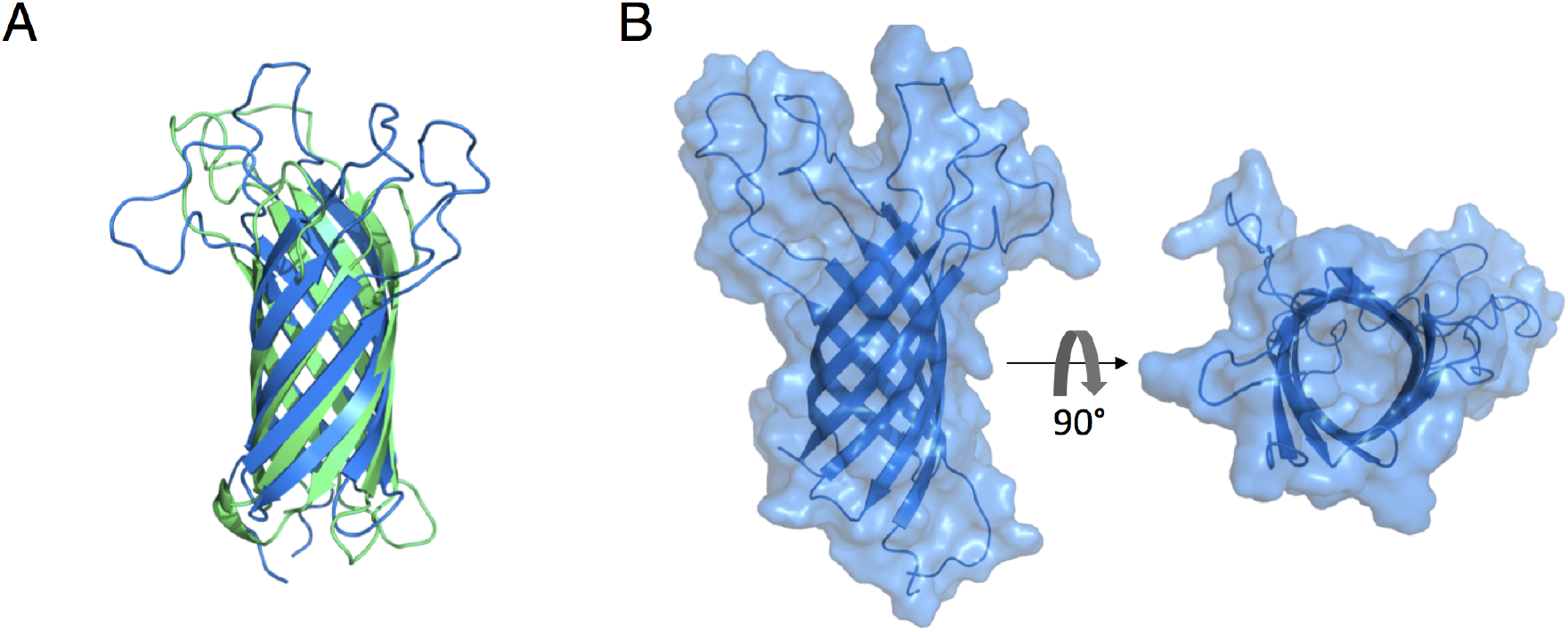
The 8-stranded β-barrels of OmpA and OmpX are structurally similar and do not display an open channel through the membrane. (**A**) Superimposition of the structures of the membrane-embedded β-barrels of OmpA (blue; PDB: 1QJP; residues 1-170) and OmpX (green; PDB: 2M07). The two β-barrels, which have no sequence homology, can be superimposed with an RMSD of 2.49 Å. (**B**) Surface representation of the β-barrel domain of OmpA (PDB: 1QJP; residues 1-170). The right panel is rotated 90° around the horizontal axis from the left panel. In this conformation, OmpA does not display an open channel.

**Figure 3 — figure supplement 2.**
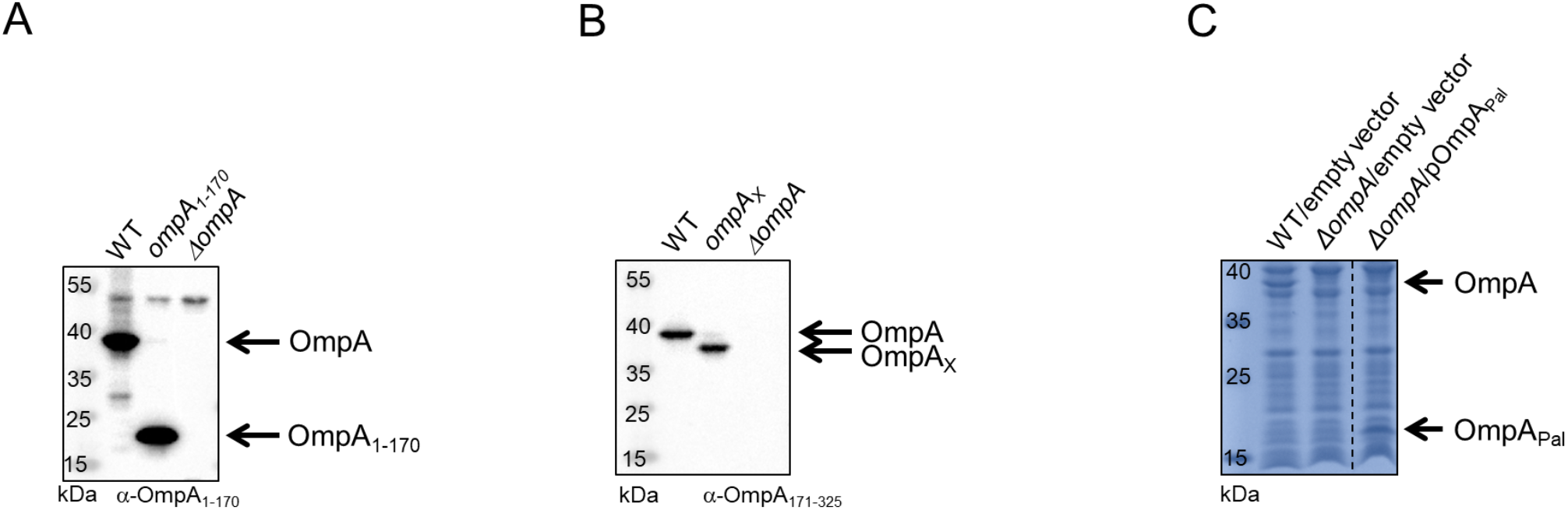
The expression levels of the OmpA variants are similar to those of wild-type OmpA. **(A,B)** Cells expressing OmpA_1-170_ **(A)** and OmpAX **(B)** from the chromosome were harvested at mid-log phase (OD_600_ = 0.4-0.6) and immunoblotted with antibodies specific for the periplasmic domain of OmpA (anti-OmpA_186-325_) or the β-barrel domain (anti-OmpA_1-170_) (Materials and Methods). **(C)** Cells harboring pKiD22 were harvested at mid-log phase (OD_600_=0.4-0.6). Expression of OmpA_Pal_ was induced with IPTG (20 μM). The membrane fraction was analyzed by SDS-PAGE and proteins stained with Coomassie Blue.

**Figure 5 — figure supplement 1:**
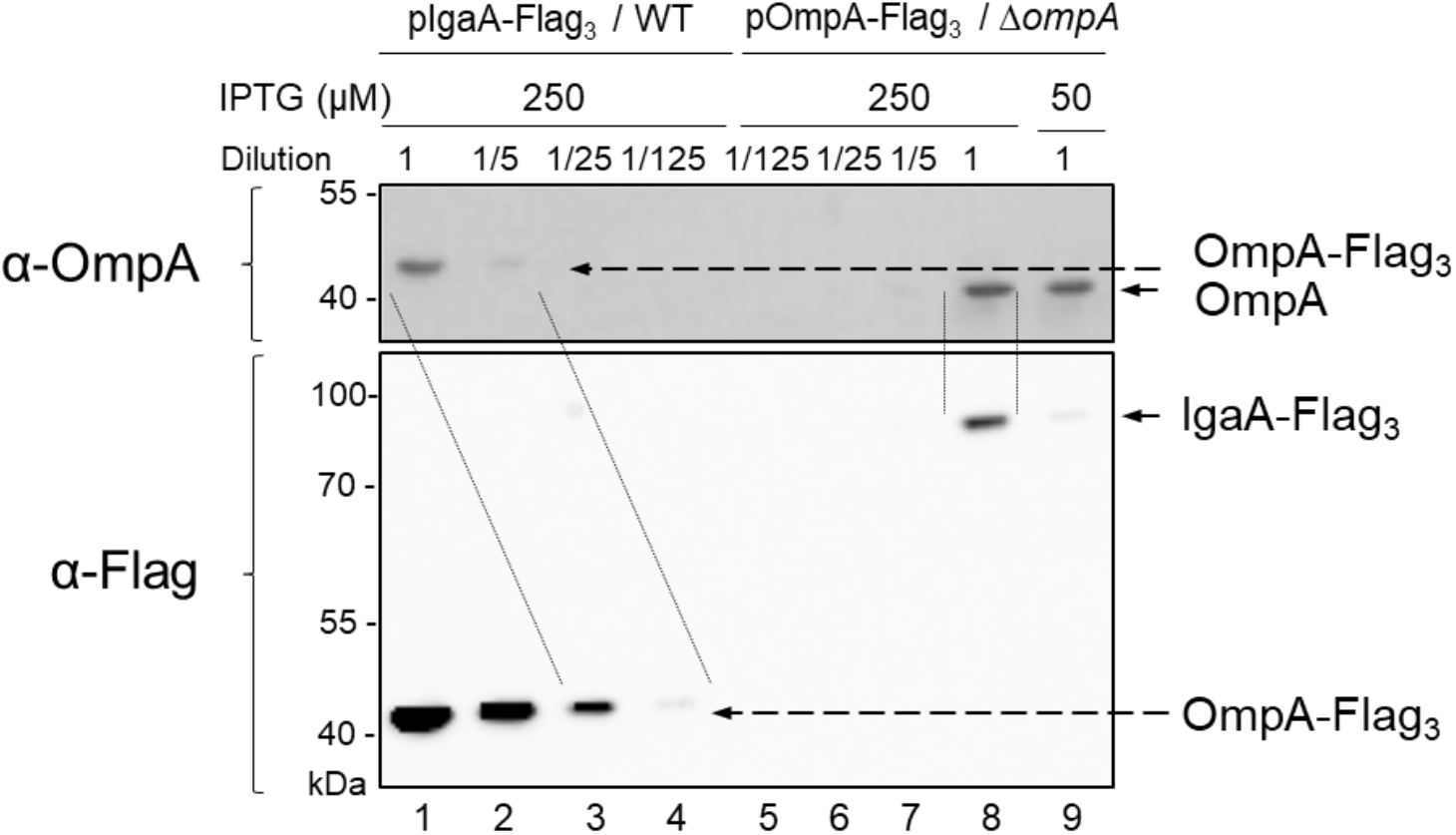
Estimation of the concentrations of IgaA and OmpA in the cell envelope. OmpA-Flag_3_ was expressed from pOmpA-Flag_3_ (pPR4) with 250 μM IPTG in *ΔompA* cells. IgaA-Flag_3_ was expressed from pIgaA-Flag_3_ (pSC237) with 50 or 250 μM IPTG in wild-type (WT) cells. Samples were serially diluted as indicated before SDS-PAGE. OmpA-Flag_3_ (lanes 1-4) and OmpA (lanes 5-9) were detected with anti-OmpA (upper panel); OmpA-Flag_3_ (lanes 1-4) and IgaA-Flag_3_ (lanes 5-9) were detected with anti-Flag (lower panel). Images are representative of experiments made in biological triplicate.

**Figure 6 — figure supplement 1:**
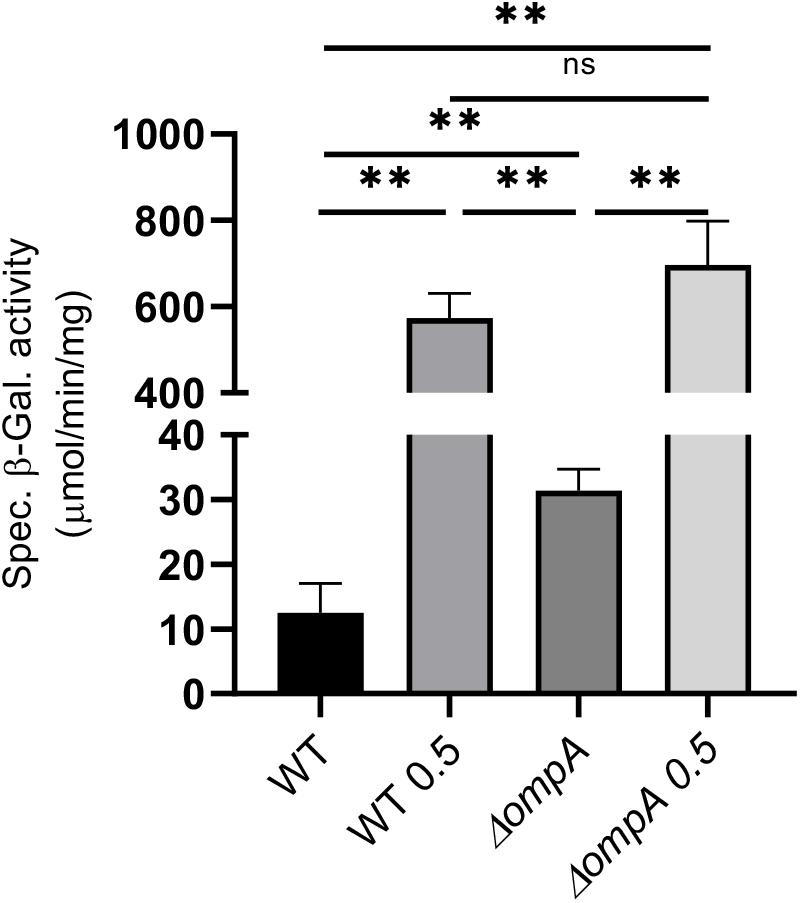
The Rcs system responds to polymyxin exposure in cells lacking OmpA. Wild-type (WT; DH300, a MG1655 derivative (Majdalani, Hernandez, and Gottesman 2002)) and *ΔompA* cells were treated with 0.5 μg/ml polymyxin B when they reached an OD_600_ of 0.4. Rcs activity was measured by a chromosomal *rprA::lacZ* fusion. Treatment with polymyxin B activated Rcs both in the WT and in the *ΔompA* mutant. Mean (n=3) and standard deviation (error bars) are shown. Differences were evaluated with Student’s *t* test (ns, not significant; **p<0.01).

**Figure 6 — figure supplement 2:**
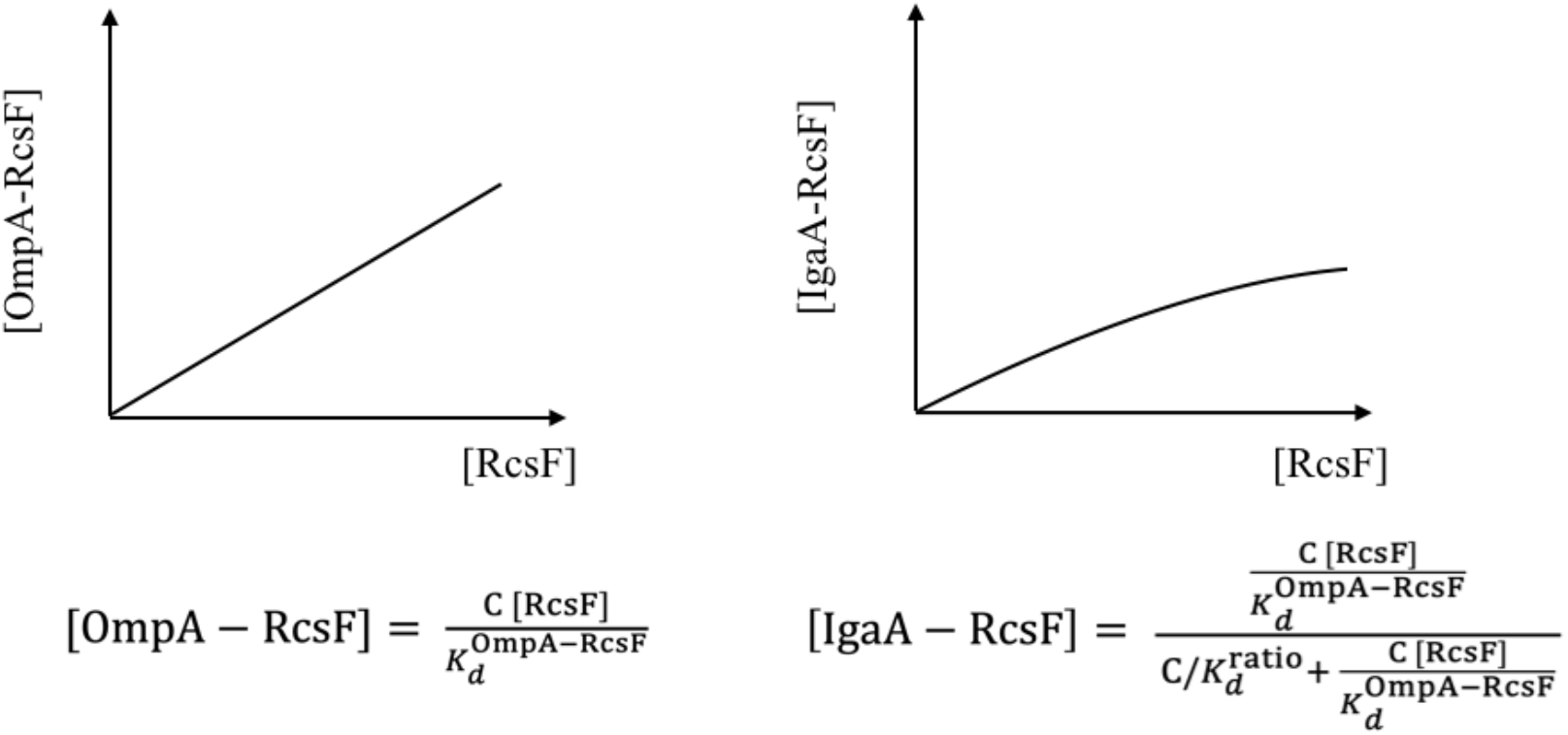
OmpA functions as a buffer for RcsF. The concentration of OmpA [OmpA] is high (1000 μM) and can therefore be considered constant ([OmpA]=C in the equations above)). Left panel: the concentration of the OmpA-RcsF complex [OmpA-RcsF] increases linearly with the concentration of RcsF in the periplasm. Right panel: the concentration of IgaA-RcsF [IgaA-RcsF] increases as a function of [RcsF] in the periplasm, but, in this case, the increase is decelerated and follows a hyperbolic curve. Thus, OmpA functions as a buffer for RcsF. See the Materials and Methods for more information.

## Supplementary Tables

**Supplementary Table 1.** Strains used in this study.

| Strains | Relevant Genotype or Features | Source or notes |
| --- | --- | --- |
| XL1-Blue | *endA1 gyrA96(nal<sup>R</sup>) thi-1 recA1 relA1 lac glnV4'*<br>F' [::Tn10 *proAB<sup>+</sup> lacI<sup>q</sup> Δ(lacZ)M15*] *hsdR17(r<sub>K</sub><sup>-</sup> m<sub>K</sub><sup>+</sup>)* | Stratagene |
| BL21 (DE3) | *E. coli* str. B F<sup>-</sup> *ompT gal dcm lon hsdS<sub>B</sub>(r<sub>B</sub><sup>-</sup> m<sub>B</sub><sup>-</sup>) λ(DE3*<br>*[lacI lacUV5-T7p07 ind1 sam7 nin5]) [malB<sup>+</sup>]<sub>K-12</sub>(λ<sup>S</sup>)* | Novagen |
| SHuffle T7 | *fhuA2 lacZ::T7 gene1 [lon] ompT ahpC gal*<br>*λatt::pNEB3-r1-cDsbC (SpecR, lacIq) ΔtrxB sulA11*<br>*R(mcr-73::miniT-0--TetS)2 [dcm] R(zgb-210::Tn- --*<br>*TetS) endA1 Δgor Δ(mcrC-mrr)114::IS10* | New England Biolabs<br>(NEB) |
| DH300 | *rprA-lacZ* MG1655 (*argF-lac*)U169 | (Majdalani,<br>Hernandez, and<br>Gottesman 2002) |
| Keio collection<br>single mutants | *ΔrcsF::kan, ΔompA::kan* | (Baba et al. 2006) |
| PL358 | DH300 *ΔrcsF* | (Cho et al. 2014) |
| SEN588 | DH300 *ΔompA::kan* | (Cho et al. 2014) |
| PR46 | DH300 *ΔompA* | this study |
| SEN589 | PL358 *ΔompA::kan* | (Cho et al. 2014) |
| SEN524 | PL358 *ΔompR::kan* | this study |
| SEN550 | DH300 pSC237 | this study |
| SEN860 | DH300 *ΔompA::cat, sacB* | this study |
| SEN861 | PL358 *ΔompA::cat, sacB* | this study |
| SEN1194 | SEN860 *ompA-6×His* | this study |
| SEN905 | SEN861 *ompA-6×His* | this study |
| SEN900 | SEN860 *ompA<sub>TH189</sub>-6×His* | this study |
| SEN901 | SEN860 *ompA<sub>TH243</sub>-6×His* | this study |
| SEN962 | SEN900 *ΔrcsF::kan* | this study |
| SEN964 | SEN901 *ΔrcsF::kan* | this study |
| SEN856 | DH300 *ΔompA<sub>171-325</sub>::kan* | this study |
| SEN857 | PL358 *ΔompA<sub>171-325</sub>::kan* | this study |
| SEN858 | DH300 *ΔompA<sub>171-325</sub>* | this study |
| SEN859 | PL358 *ΔompA<sub>171-325</sub>* | this study |
| SEN1089 | PL358 pSUP-Mb_DiZPK-RS and pSC253 | this study |
| SEN1091 | PL358 pSUP-Mb_DiZPK-RS and pSC253(Q79X) | this study |
| SEN1094 | PL358 pSUP-Mb_DiZPK-RS and pSC253(P116X) | this study |
| SEN1109 | PL358 pSUP-Mb_DiZPK-RS and pSC253(K98X) | this study |
| SEN1113 | PL358 pSUP-Mb_DiZPK-RS and pSC253(E110X) | this study |
| SEN1115 | PL358 pSUP-Mb_DiZPK-RS and pSC253(Q121X) | this study |
| SEN1129 | PL358 pSUP-Mb_DiZPK-RS and pSC253(R21X) | this study |
| SEN1130 | PL358 pSUP-Mb_DiZPK-RS and pSC253(Q28X) | this study |
| SEN1131 | PL358 pSUP-Mb_DiZPK-RS and pSC253(Q33X) | this study |
| SEN1134 | PL358 pSUP-Mb_DiZPK-RS and pSC253(R45X) | this study |
| SEN1136 | PL358 pSUP-Mb_DiZPK-RS and pSC253(N54X) | this study |
| SEN1141 | PL358 pSUP-Mb_DiZPK-RS and pSC253(R89X) | this study |
| SEN1275 | SEN589 pSUP-Mb_DiZPK-RS and pSC253(R89X) | this study |
| SEN1276 | SEN524 pSUP-Mb_DiZPK-RS and pSC253(R89X) | this study |
| SEN1689 | SEN589 pSUP-Mb_DiZPK-RS and pSC253(K98X) | this study |
| SEN1690 | SEN589 pSUP-Mb_DiZPK-RS and pSC253(E110X) | this study |
| SEN1378 | PR46 pSUP-Mb_DiZPK-RS and pPR21 | this study |
| SEN1381 | PR46 pSUP-Mb_DiZPK-RS and pPR21(R242X) | this study |
| SEN1552 | DH300 pSC231 | this study |
| SEN1562 | PR46 pSC237 | this study |
| SEN1565 | PL358 pSC237 | this study |
| PR44 | PR46 pPR4 | this study |
| JLE60 | SEN860 $\Delta ompA::ompX-ompA_{171-325}$ | this study |
| JLE61 | SEN861 $\Delta ompA::ompX-ompA_{171-325}$ | this study |
| KiD003 | BL21(DE3) pKiD5 | this study |
| KiD083 | PR46 pSUP-Mb_DiZPK-RS and pPR21(D246X) | this study |
| KiD084 | PR46 pSUP-Mb_DiZPK-RS and pPR21(Y248X) | this study |
| KiD123 | DH300 pKiD22 | this study |
| KiD124 | PR46 pKiD22 | this study |

**Supplementary Table 2.**
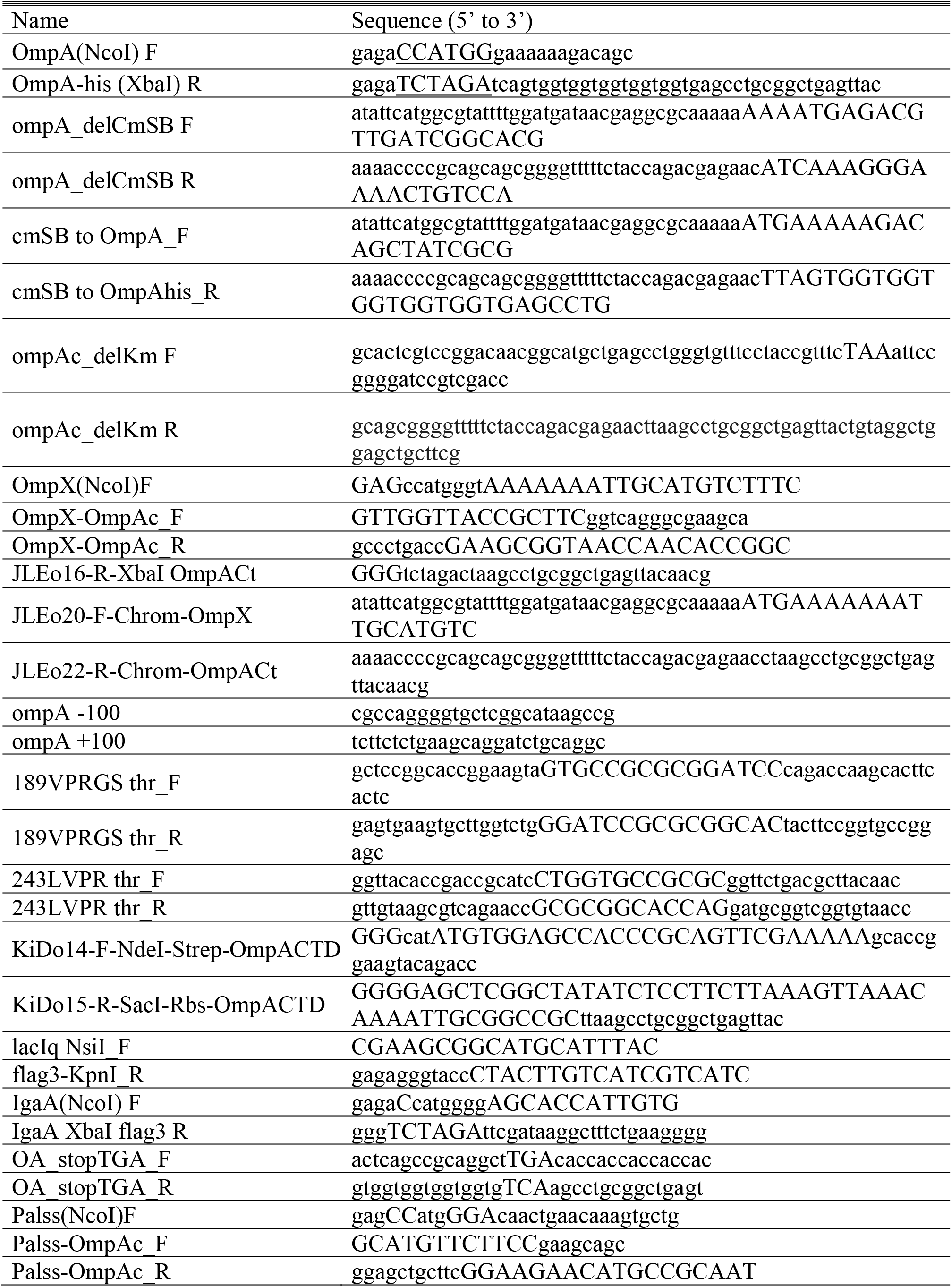
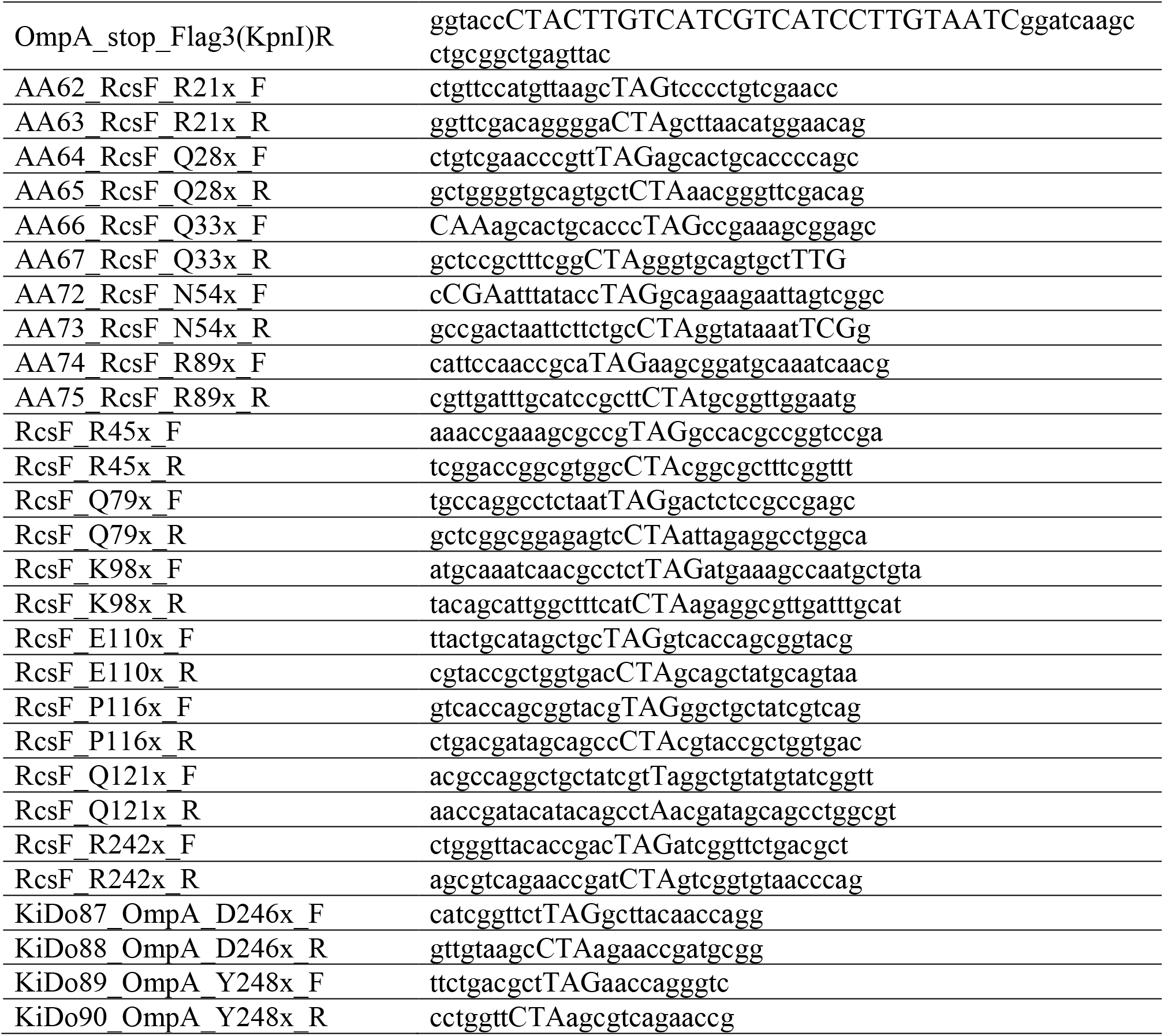
Primers used in this study.

**Supplementary Table 3.** Plasmids used in this study.

| Plasmids | Features | Source or notes |
| --- | --- | --- |
| pDSW204 | IPTG-regulated modified $P_{trc}$, ampicillin | (Weiss et al. 1999) |
| pAM238 | IPTG-regulated $P_{lac}$, pSC101-based, spectinomycin | (Gil and Bouche 1991) |
| pBAD18 | Arabinose regulation, ampicillin | (Guzman et al. 1995) |
| pSup-Mb-DIZPK-RS | PylRS, tRNA<sup>Pyl</sup><sub>CUA</sub> opt, p15A origin, chloramphenicol | (Zhang et al. 2011) |
| pSIM5-tet | pSC101-based, $repA^{ts}$, tetracycline | (Koskiniemi et al. 2011) |
| pCP20 | $FLP^+$, $\lambda$ cI857<sup>+</sup>, $\lambda$ $P_{PR}$ Rep<sup>ts</sup>, ampicillin, chloramphenicol | (Cherepanov and Wackernagel 1995) |
| pSC202 | pAM238 with RcsF | (Cho et al. 2014) |
| pSC253 | pBAD18 with RcsF | this study |
| pPR11 | pDSW204 with OmpA-6×His | this study |
| pPR11<sub>189thrombin</sub> | pDSW204 with OmpA(189-Val-Val-Pro-Arg-Gly-Ser-Gln-190)-6×His | this study |
| pPR11<sub>243thrombin</sub> | pDSW204 with OmpA(243-Ile-Leu-Val-Pro-Arg-Gly-244)-6×His | this study |
| pJLE17-OmpX-OmpA<sub>171-325</sub> | pDSW204 with OmpA<sub>X</sub> | this study |
| pKiD5 | pET21a with strep-OmpA<sub>186-325</sub> | this study |
| pMER77 | pDSW204 with Flag<sub>3</sub> | (Hemmis et al. 2011) |
| pSC231 | pAM238 with LacI<sup>q</sup>, the modified $P_{trc}$, and Flag<sub>3</sub> | this study |
| pPR4 | pSC231 with OmpA | this study |
| pSC237 | pSC231 with IgaA | this study |
| pPR21 | pDSW204 with OmpA | this study |
| pKiD22 | pDSW204 with OmpA<sub>Pal</sub> | this study |

## References

Abraham, Mark James, Teemu Murtola, Roland Schulz, Szilárd Páll, Jeremy C. Smith, Berk Hess, and Erik Lindahl. 2015. “GROMACS: High performance molecular simulations through multi-level parallelism from laptops to supercomputers.” SoftwareX 1-2:19–25. doi: https://doi.org/10.1016/j.softx.2015.06.001.

Baba, T., T. Ara, M. Hasegawa, Y. Takai, Y. Okumura, M. Baba, K. A. Datsenko, M. Tomita, B. L. Wanner, and H. Mori. 2006. “Construction of Escherichia coli K-12 in-frame, singlegene knockout mutants: the Keio collection.” Mol Syst Biol 2:2006 0008. doi: msb4100050 [pii] 10.1038/msb4100050.

Basle, A., G. Rummel, P. Storici, J. P. Rosenbusch, and T. Schirmer. 2006. “Crystal structure of osmoporin OmpC from E. coli at 2.0 A.” J Mol Biol 362 (5):933–42. doi: 10.1016/j.jmb.2006.08.002.

Cascales, E., A. Bernadac, M. Gavioli, J. C. Lazzaroni, and R. Lloubes. 2002. “Pal lipoprotein of Escherichia coli plays a major role in outer membrane integrity.” J Bacteriol 184 (3):754–9.

Chai, T. J., and J. Foulds. 1977. “Purification of protein A, an outer membrane component missing in Escherichia coli K-12 ompA mutants.” Biochim Biophys Acta 493 (1):210–5. doi: 10.1016/0005-2795(77)90274-4.

Cherepanov, P. P., and W. Wackernagel. 1995. “Gene disruption in Escherichia coli: TcR and KmR cassettes with the option of Flp-catalyzed excision of the antibiotic-resistance determinant.” Gene 158 (1):9–14.

Chin, J. W., A. B. Martin, D. S. King, L. Wang, and P. G. Schultz. 2002. “Addition of a photocrosslinking amino acid to the genetic code of Escherichiacoli.” Proc Natl Acad Sci US A 99 (17):11020–4. doi: 10.1073/pnas.172226299.

Cho, S. H., J. Szewczyk, C. Pesavento, M. Zietek, M. Banzhaf, P. Roszczenko, A. Asmar, G. Laloux, A. K. Hov, P. Leverrier, C. Van der Henst, D. Vertommen, A. Typas, and J. F. Collet. 2014. “Detecting Envelope Stress by Monitoring beta-Barrel Assembly.” Cell 159 (7):1652–64. doi: 10.1016/j.cell.2014.11.045.

Chubiz, Lon M, and Christopher V Rao. 2011. “Role of the mar-sox-rob regulon in regulating outer membrane porin expression.” Journal of bacteriology 193 (9):2252–2260.

Datsenko, K. A., and B. L. Wanner. 2000. “One-step inactivation of chromosomal genes in Escherichia coli K-12 using PCR products.” Proc Natl Acad Sci U S A 97 (12):6640–5. doi: 10.1073/pnas.120163297 120163297 [pii].

De Mot, R., and J. Vanderleyden. 1994. “The C-terminal sequence conservation between OmpA-related outer membrane proteins and MotB suggests a common function in both grampositive and gram-negative bacteria, possibly in the interaction of these domains with peptidoglycan.” Mol Microbiol 12 (2):333–4. doi: 10.1111/j.1365-2958.1994.tb01021.x.

Delhaye, Antoine, Jean-Francois Collet, and Géraldine Laloux. 2019. “A fly on the wall: how stress response systems can sense and respond to damage to peptidoglycan.” Frontiers in Cellular and Infection Microbiology 9.

Dominguez-Bernal, G., M. G. Pucciarelli, F. Ramos-Morales, M. Garcia-Quintanilla, D. A. Cano, J. Casadesus, and F. Garcia-del Portillo. 2004. “Repression of the RcsC-YojN-RcsB phosphorelay by the IgaA protein is a requisite for Salmonella virulence.” Mol Microbiol 53 (5):1437–49. doi: 10.1111/j.1365-2958.2004.04213.x.

Egan, A. J. F., R. Maya-Martinez, I. Ayala, C. M. Bougault, M. Banzhaf, E. Breukink, W. Vollmer, and J. P. Simorre. 2018. “Induced conformational changes activate the peptidoglycan synthase PBP1B.” Mol Microbiol 110 (3):335–356. doi: 10.1111/mmi.14082.

Egan, Alexander JF, Jeff Errington, and Waldemar Vollmer. 2020. “Regulation of peptidoglycan synthesis and remodelling.” Nature Reviews Microbiology:1–15.

Farris, C., S. Sanowar, M. W. Bader, R. Pfuetzner, and S. I. Miller. 2010. “Antimicrobial peptides activate the Rcs regulon through the outer membrane lipoprotein RcsF.” J Bacteriol 192 (19):4894–903. doi: JB.00505-10 [pii] 10.1128/JB.00505-10.

Favier, A., and B. Brutscher. 2011. “Recovering lost magnetization: polarization enhancement in biomolecular NMR.” J Biomol NMR 49 (1):9–15. doi: 10.1007/s10858-010-9461-5.

Gennaris, A., B. Ezraty, C. Henry, R. Agrebi, A. Vergnes, E. Oheix, J. Bos, P. Leverrier, L. Espinosa, J. Szewczyk, D. Vertommen, O. Iranzo, J. F. Collet, and F. Barras. 2015. “Repairing oxidized proteins in the bacterial envelope using respiratory chain electrons.” Nature 528 (7582):409–12. doi: 10.1038/nature15764.

Gil, D., and J. P. Bouche. 1991. “ColE1-type vectors with fully repressible replication.” Gene 105 (1):17–22.

Guzman, L. M., D. Belin, M. J. Carson, and J. Beckwith. 1995. “Tight regulation, modulation, and high-level expression by vectors containing the arabinose PBAD promoter.” J Bacteriol 177 (14):4121–30.

Hall, MICHAEL N, and THOMAS J Silhavy. 1979. “Transcriptional regulation of Escherichia coli K-12 major outer membrane protein 1b.” Journal of bacteriology 140 (2):342–350.

Hemmis, C. W., M. Berkmen, M. Eser, and J. F. Schildbach. 2011. “TrbB from conjugative plasmid F is a structurally distinct disulfide isomerase that requires DsbD for redox state maintenance.” J Bacteriol 193 (18):4588–97. doi: 10.1128/JB.00351-11.

Housden, N. G., J. T. Hopper, N. Lukoyanova, D. Rodriguez-Larrea, J. A. Wojdyla, A. Klein, R. Kaminska, H. Bayley, H. R. Saibil, C. V. Robinson, and C. Kleanthous. 2013. “Intrinsically disordered protein threads through the bacterial outer-membrane porin OmpF.” Science 340 (6140):1570–4. doi: 10.1126/science.1237864.

Hussain, S., and H. D. Bernstein. 2018. “The Bam complex catalyzes efficient insertion of bacterial outer membrane proteins into membrane vesicles of variable lipid composition.” J Biol Chem 293 (8):2959–2973. doi: 10.1074/jbc.RA117.000349.

Hussein, N. A., S. H. Cho, G. Laloux, R. Siam, and J. F. Collet. 2018. “Distinct domains of Escherichia coli IgaA connect envelope stress sensing and down-regulation of the Rcs phosphorelay across subcellular compartments.” PLoS Genet 14 (5):e1007398. doi: 10.1371/journal.pgen.1007398.

Ishida, H., A. Garcia-Herrero, and H. J. Vogel. 2014. “The periplasmic domain of Escherichia coli outer membrane protein A can undergo a localized temperature dependent structural transition.” Biochim Biophys Acta 1838 (12):3014–24. doi: 10.1016/j.bbamem.2014.08.008.

Jorgensen, William L., David S. Maxwell, and Julian Tirado-Rives. 1996. “Development and Testing of the OPLS All-Atom Force Field on Conformational Energetics and Properties of Organic Liquids.” Journal of the American Chemical Society 118 (45): 11225–11236. doi: 10.1021/ja9621760.

Konovalova, A., A. M. Mitchell, and T. J. Silhavy. 2016. “A lipoprotein/beta-barrel complex monitors lipopolysaccharide integrity transducing information across the outer membrane.” Elife 5. doi: 10.7554/eLife.15276.

Konovalova, A., D. H. Perlman, C. E. Cowles, and T. J. Silhavy. 2014. “Transmembrane domain of surface-exposed outer membrane lipoprotein RcsF is threaded through the lumen of beta-barrel proteins.” Proc Natl Acad Sci U S A 111 (41):E4350–8. doi: 10.1073/pnas.1417138111.

Konovalova, Anna, and Thomas J Silhavy. 2015. “Outer membrane lipoprotein biogenesis: Lol is not the end.” Philosophical Transactions of the Royal Society B: Biological Sciences 370 (1679):20150030.

Koskiniemi, S., M. Pranting, E. Gullberg, J. Nasvall, and D. I. Andersson. 2011. “Activation of cryptic aminoglycoside resistance in Salmonella enterica.” Mol Microbiol 80 (6):1464–78. doi: 10.1111/j.1365-2958.2011.07657.x.

Laloux, G., and J. F. Collet. 2017. ““Major Tom to ground control: how lipoproteins communicate extra-cytoplasmic stress to the decision center of the cell”.” J Bacteriol. doi: 10.1128/JB.00216-17.

Laubacher, M. E., and S. E. Ades. 2008. “The Rcs phosphorelay is a cell envelope stress response activated by peptidoglycan stress and contributes to intrinsic antibiotic resistance.” J Bacteriol 190 (6):2065–74. doi: JB.01740-07 [pii] 10.1128/JB.01740-07.

Leverrier, P., J. P. Declercq, K. Denoncin, D. Vertommen, A. Hiniker, S. H. Cho, and J. F. Collet. 2011. “Crystal structure of the outer membrane protein RcsF, a new substrate for the periplasmic protein-disulfide isomerase DsbC.” J Biol Chem 286 (19):16734–42. doi: 10.1074/jbc.M111.224865.

Li G-W, Burkhardt DH, Gross CA, Weissman JS. 2014. “Quantifying absolute protein synthesis rates reveals principles underlying allocation of cellular resources.” Cell 157:624–635.

Li, G. W., D. Burkhardt, C. Gross, and J. S. Weissman. 2014. “Quantifying absolute protein synthesis rates reveals principles underlying allocation of cellular resources.” Cell 157 (3):624–35. doi: 10.1016/j.cell.2014.02.033.

MacRitchie, D. M., D. R. Buelow, N. L. Price, and T. L. Raivio. 2008. “Two-component signaling and gram negative envelope stress response systems.” Adv Exp Med Biol 631:80–110. doi: 10.1007/978-0-387-78885-2_6.

Majdalani, N., D. Hernandez, and S. Gottesman. 2002. “Regulation and mode of action of the second small RNA activator of RpoS translation, RprA.” Mol Microbiol 46 (3):813–26.

Neidhardt, F. C., P. L. Bloch, and D. F. Smith. 1974. “Culture medium for enterobacteria.” J Bacteriol 119 (3):736–47.

Noinaj, N., J. C. Gumbart, and S. K. Buchanan. 2017. “The beta-barrel assembly machinery in motion.” Nat Rev Microbiol 15 (4):197–204. doi: 10.1038/nrmicro.2016.191.

Okuda, S., D. J. Sherman, T. J. Silhavy, N. Ruiz, and D. Kahne. 2016. “Lipopolysaccharide transport and assembly at the outer membrane: the PEZ model.” Nat Rev Microbiol 14 (6):337–45. doi: 10.1038/nrmicro.2016.25.

Park, J. S., W. C. Lee, K. J. Yeo, K. S. Ryu, M. Kumarasiri, D. Hesek, M. Lee, S. Mobashery, J. H. Song, S. I. Kim, J. C. Lee, C. Cheong, Y. H. Jeon, and H. Y. Kim. 2012. “Mechanism of anchoring of OmpA protein to the cell wall peptidoglycan of the gram-negative bacterial outer membrane.” FASEB J 26 (1):219–28. doi: 10.1096/fj.11-188425.

Pautsch, A., and G. E. Schulz. 1998. “Structure of the outer membrane protein A transmembrane domain.” Nat Struct Biol 5 (11):1013–7. doi: 10.1038/2983.

Radhakrishnan, S. K., S. Pritchard, and P. H. Viollier. 2010. “Coupling prokaryotic cell fate and division control with a bifunctional and oscillating oxidoreductase homolog.” Dev Cell 18 (1):90–101. doi: 10.1016/j.devcel.2009.10.024.

Reusch, R. N. 2012. “Insights into the structure and assembly of Escherichia coli outer membrane protein A.” Febs j 279 (6):894–909. doi: 10.1111/j.1742-4658.2012.08484.x.

Rodriguez-Alonso, R., J. Letoquart, V. S. Nguyen, G. Louis, A. N. Calabrese, B. I. Iorga, S. E. Radford, S. H. Cho, H. Remaut, and J. F. Collet. 2020. “Structural insight into the formation of lipoprotein-beta-barrel complexes.” Nat Chem Biol. doi: 10.1038/s41589-020-0575-0.

Rogov, V. V., N. Y. Rogova, F. Bernhard, F. Lohr, and V. Dotsch. 2011. “A disulfide bridge network within the soluble periplasmic domain determines structure and function of the outer membrane protein RCSF.” J Biol Chem 286 (21):18775–83. doi: 10.1074/jbc.M111.230185.

Ruiz, N., and T. J. Silhavy. 2005. “Sensing external stress: watchdogs of the Escherichia coli cell envelope.” Curr Opin Microbiol 8 (2): 122–6.

Silhavy, T. J., D. Kahne, and S. Walker. 2010. “The bacterial cell envelope.” Cold Spring Harb Perspect Biol 2 (5):a000414. doi: 10.1101/cshperspect.a000414.

Singh, S. P., Y. U. Williams, S. Miller, and H. Nikaido. 2003. “The C-terminal domain of Salmonella enterica serovar typhimurium OmpA is an immunodominant antigen in mice but appears to be only partially exposed on the bacterial cell surface.” Infect Immun 71 (7):3937–46. doi: 10.1128/iai.71.7.3937-3946.2003.

Stathopoulos, C. 1996. “An alternative topological model for Escherichia coli OmpA.” Protein Sci 5 (1):170–3. doi: 10.1002/pro.5560050122.

Stock, J. B., B. Rauch, and S. Roseman. 1977. “Periplasmic space in Salmonella typhimurium and Escherichia coli.” J Biol Chem 252 (21):7850–61.

Szewczyk, J., and J. F. Collet. 2016. “The Journey of Lipoproteins Through the Cell: One Birthplace, Multiple Destinations.” Adv Microb Physiol 69:1–50. doi: 10.1016/bs.ampbs.2016.07.003.

Takeda, S., Y. Fujisawa, M. Matsubara, H. Aiba, and T. Mizuno. 2001. “A novel feature of the multistep phosphorelay in Escherichia coli: a revised model of the RcsC --> YojN --> RcsB signalling pathway implicated in capsular synthesis and swarming behaviour.” Mol Microbiol 40 (2):440–50. doi: mmi2393 [pii].

Thomason, L. C., J. A. Sawitzke, X. Li, N. Costantino, and D. L. Court. 2014. “Recombineering: genetic engineering in bacteria using homologous recombination.” Curr Protoc Mol Biol 106:1 16 1–39. doi: 10.1002/0471142727.mb0116s106.

Typas, A., M. Banzhaf, C. A. Gross, and W. Vollmer. 2012. “From the regulation of peptidoglycan synthesis to bacterial growth and morphology.” Nat Rev Microbiol 10 (2):123–36. doi: 10.1038/nrmicro2677.

van Zundert, G. C. P., Jpglm Rodrigues, M. Trellet, C. Schmitz, P. L. Kastritis, E. Karaca, A. S. J. Melquiond, M. van Dijk, S. J. de Vries, and A M J J Bonvin. 2016. “The HADDOCK2.2 Web Server: User-Friendly Integrative Modeling of Biomolecular Complexes.” J Mol Biol 428 (4):720–725. doi: 10.1016/j.jmb.2015.09.014.

Vaynberg, J., and J. Qin. 2006. “Weak protein-protein interactions as probed by NMR spectroscopy.” Trends Biotechnol 24 (1):22–7. doi: 10.1016/j.tibtech.2005.09.006.

Wall, E., N. Majdalani, and S. Gottesman. 2018. “The Complex Rcs Regulatory Cascade.” Annu Rev Microbiol 72:111–139. doi: 10.1146/annurev-micro-090817-062640.

Wall, Erin A, Nadim Majdalani, and Susan Gottesman. 2020. “Negative regulation of the Rcs phosphorelay via IgaA contact with the RcsD phosphotransfer protein.” bioRxiv.

Wassenaar, Tsjerk A., Marc van Dijk, Nuno Loureiro-Ferreira, Gijs van der Schot, Sjoerd J. de Vries, Christophe Schmitz, Johan van der Zwan, Rolf Boelens, Andrea Giachetti, Lucio Ferella, Antonio Rosato, Ivano Bertini, Torsten Herrmann, Hendrik R. A. Jonker, Anurag Bagaria, Victor Jaravine, Peter Güntert, Harald Schwalbe, Wim F. Vranken, Jurgen F. Doreleijers, Gert Vriend, Geerten W. Vuister, Daniel Franke, Alexey Kikhney, Dmitri I. Svergun, Rasmus H. Fogh, John Ionides, Ernest D. Laue, Chris Spronk, Simonas Jurkša, Marco Verlato, Simone Badoer, Stefano Dal Pra, Mirco Mazzucato, Eric Frizziero, and Alexandre M. J. J. Bonvin. 2012. “WeNMR: Structural Biology on the Grid.” Journal of Grid Computing 10 (4):743–767. doi: 10.1007/s10723-012-9246-z.

Weiss, D. S., J. C. Chen, J. M. Ghigo, D. Boyd, and J. Beckwith. 1999. “Localization of FtsI (PBP3) to the septal ring requires its membrane anchor, the Z ring, FtsA, FtsQ, and FtsL.” J Bacteriol 181 (2):508–20.

Yamashita, E., M. V. Zhalnina, S. D. Zakharov, O. Sharma, and W. A. Cramer. 2008. “Crystal structures of the OmpF porin: function in a colicin translocon.” EMBO J 27 (15):2171–80. doi: 10.1038/emboj.2008.137.

Yu, D., H. M. Ellis, E. C. Lee, N. A. Jenkins, N. G. Copeland, and D. L. Court. 2000. “An efficient recombination system for chromosome engineering in Escherichia coli.” Proc Natl Acad Sci USA 97 (11):5978–83. doi: 10.1073/pnas.100127597.

Zhang, M., S. Lin, X. Song, J. Liu, Y. Fu, X. Ge, X. Fu, Z. Chang, and P. R. Chen. 2011. “A genetically incorporated crosslinker reveals chaperone cooperation in acid resistance.” Nat Chem Biol 7 (10):671–7. doi: 10.1038/nchembio.644.

Zhang, X., and H. Bremer. 1995. “Control of the Escherichia coli rrnB P1 promoter strength by ppGpp.” J Biol Chem 270 (19):11181–9. doi: 10.1074/jbc.270.19.11181.

